# Normalization accounts for temporal dynamics in human somatosensory cortex

**DOI:** 10.64898/2026.05.10.724117

**Authors:** Ilona M. Bloem, Luhe Li, Stephanie Badde, Iris I.A. Groen, Wouter Schellekens, Nick Ramsey, Adeen Flinker, Orrin Devinsky, Sasha Devore, Werner Doyle, Patricia Dugan, Daniel Friedman, Natalia Petridou, Michael S. Landy, Jonathan Winawer

**Author notes:** Correspondence should be addressed to Ilona Bloem. These authors provided equal contribution (Ilona M. Bloem, Luhe Li). The authors declare no competing financial interests.

## Abstract

Sensory processing is fundamentally shaped by stimulation history. For example, in visual cortex, neural responses are reduced for repeated or sustained stimuli (adaptation). These phenomena are well characterized and effectively modeled by divisive normalization. We asked whether these same computational principles govern somatosensory processing. We used fMRI (6 participants) and intracranial electroencephalography (iEEG, 2 participants) to measure responses to time-varying vibrotactile stimuli in human somatosensory cortex. Stimuli consisted of single- and paired-pulses with durations and interstimulus intervals ranging from 0.05 to 1.2 s. We extracted BOLD time courses to capture neural response amplitudes, and high-frequency iEEG broadband envelopes to capture fast neural dynamics. In both experiments, we observed pronounced sub-additive temporal summation. Responses to longer or repeated stimuli were consistently lower than predicted by linear integration. Computational modeling revealed that divisive normalization models outperformed linear models in cross-validated accuracy across both datasets. These results demonstrate that somatosensory temporal dynamics closely mirror those in the visual system. Our findings suggest that the nervous system employs similar computational principles across modalities to encode sensory information across time.

**Significance statement:** How the brain integrates sensory information over time is a fundamental question in neuroscience. While nonlinear temporal integration is well documented in visual cortex, it has not been extensively mapped in the human somatosensory system. By combining fMRI with intracranial EEG in humans, we demonstrate that somatosensory responses to tactile stimulation exhibit subadditive temporal summation. This nonlinearity is accurately captured by a divisive normalization model, matching observations in the visual system. Our results suggest that normalization is a canonical computation shared across different modalities to manage temporal dynamics, providing a unified framework for understanding how the brain encodes dynamic sensory stimuli.

## 1. Introduction

Perceptual systems integrate sensory inputs across space and time. Despite vast differences across sensory modalities in the spatiotemporal properties of their inputs, as well as their transduction mechanisms and neural pathways, there are some principles that have been proposed to generalize across senses. One such principle is divisive normalization, in which a neural response is divided by the pooled activity of a surrounding population (Heeger, 1992). This operation provides a flexible mechanism for gain control, context dependence, and perceptual invariance (Carandini & Heeger, 2012), supporting robust coding under variable stimulus conditions. Normalization has been proposed as a unifying, canonical computation across sensory systems, including vision, audition, olfaction, and somatosensation (Brouwer et al., 2015; Carandini & Heeger, 2012; Olsen et al., 2010; Willmore & King, 2023), as well as higher-order processes such as attention (Bloem & Ling, 2019; Denison, 2024; Lee & Maunsell, 2009; Reynolds & Heeger, 2009) and decision-making (Keung et al., 2020; Louie et al., 2013).

In the somatosensory cortex, normalization has primarily been studied in the spatial domain. Neurons in primary somatosensory cortex are somatotopically tuned, responding selectively to touch on specific locations on the skin (Penfield & Rasmussen, 1950; Schellekens et al., 2021; Tommerdahl et al., 2010). Studies have shown that tactile responses from different skin locations are combined in a nonlinear manner (Arbuckle et al., 2022; Badde et al., 2014; DiCarlo et al., 1998; Hyvärinen & Poranen, 1978; Reed et al., 2008; Schellekens et al., 2021). When multiple fingers are stimulated, responses are combined sub-additively consistent with cross-digit suppression (Arbuckle et al., 2022; Brouwer et al., 2015). These results are well explained by models that implement divisive normalization (Brouwer et al., 2015; Carandini & Heeger, 2012). Thus, normalization appears to support spatial integration across the fingers. However, it is not yet clear whether similar computations underlie how the somatosensory system integrates information over time.

Temporal summation in visual cortex is nonlinear (Tolhurst et al., 1980), a finding that has been linked to divisive normalization (Heeger, 1992; Mikaelian & Simoncelli, 2001). Neural responses to prolonged and repeated visual stimuli show clear sub-additive temporal dynamics (Chapman & Denison, 2025; Groen et al., 2022; Zhou et al., 2018, 2019, though see Zhou et al., 2024). Prolonged stimuli lead to sub-additive responses in that the responses to longer stimuli, summed over time, are less than the linear prediction from shorter stimuli. Repeated stimuli likewise lead to a reduced response to the second stimulus presentation, as long as the gap between the stimuli is relatively short. Divisive normalization models account for these temporal non-linearities in visual responses in both fMRI (Zhou et al., 2018) and intracranial electroencephalography (iEEG; Brands et al., 2024; Groen et al., 2022; Zhou et al., 2019).

Here, we hypothesized that the same principles apply more generally to sensory coding. Just as neural responses in visual cortex undergo sub-additive temporal summation for prolonged and repeated stimuli, we predicted that similar temporal dynamics are evident for responses in somatosensory cortex. We used fMRI and iEEG to measure responses to tactile stimuli within the human somatosensory cortex. The fMRI signal can be used to measure the extent of subadditivity in temporal summation despite the slow hemodynamics as shown in the visual system (Stigliani et al., 2019; Zhou et al., 2018), whereas the iEEG signal measures both the extent and the timing of the subadditivity (Brands et al., 2024; Groen et al., 2022; Zhou et al., 2019). For both measurement modalities, we used an event-related design, with single- and double-pulse vibrotactile stimulation, comparable to the visual studies. Deconvolved BOLD response time courses and the iEEG broadband time courses revealed clear sub-additive temporal summation. The iEEG data additionally showed that the temporal dynamics depended on response history, comparable with observations from visual studies. We found that a model with divisive normalization outperformed a linear model for both datasets. These findings support the view that normalization is a canonical computation across sensory modalities.

## 2. fMRI Methods

### Participants

Six healthy participants (all female; mean age = 28.3 years old) participated in the fMRI study. The University Committee on Activities Involving Human Subjects at New York University approved the study. All participants provided written informed consent at the start of the experiment. Participants received monetary reimbursement for their time, except for one participant who is an author of the study.

### Apparatus and stimuli

The experiment was run using Matlab (R2016b, 64-bit, Mathworks, Natick, MA, USA) in conjunction with the Psychophysics Toolbox (Brainard, 1997; Pelli, 1997), using a computer running Windows 10 Professional. Participants were placed comfortably in the scanner with their heads fixed, using padding to minimize head motion.

Participants were presented with vibrotactile stimulation to the fingertip pads of the non-dominant hand using MR-compatible stimulators with a triangular-shaped tip and a contact area of around 1 mm^2^ (plectrum piezo stimulators, Dancer Design, Ingleton, UK; see **Figure 1a**). The stimulation was delivered through custom-written Matlab code in conjunction with the National Instruments toolbox. Stimulus signals created in Matlab were converted into analog signals using an NI-9264 digital-to-analog converter output module (National Instruments, Austin, TX, USA), connected to the Windows PC and subsequently amplified (piezo amplifier, Dancer Design). Stimulators were mounted by using velcro to secure a custom mold grouping 3 stimulators together on each fingertip (a total of 15 stimulators). Foam padding was added between fingers to ensure digits would not touch each other. The participant’s hand was placed with the palm facing down on their stomach. A piece of dense foam was placed underneath the hand, preventing tactile stimulation of the stomach and minimizing hand movements.

**Figure 1.**
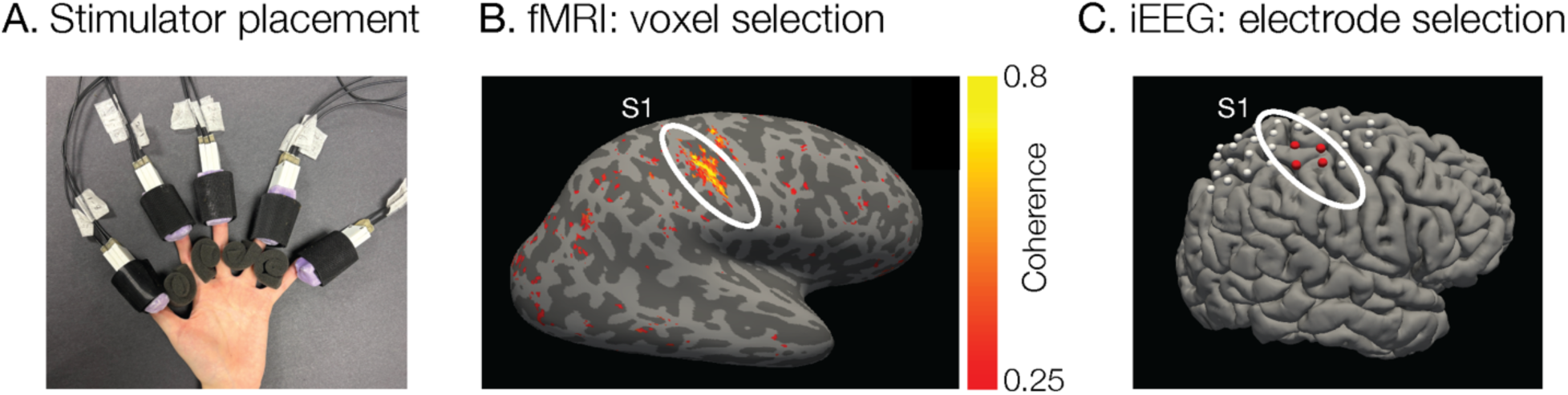
Tactile stimulation setup and regions of interest. **A.** Three MRI-compatible piezo stimulators were attached to each fingertip to deliver vibrotactile stimulation. **B.** The region of interest in the primary somatosensory cortex was functionally localized for all participants. Coherence map for an example fMRI participant obtained by a phase-encoded analysis. **C.** Intracranial EEG electrode coverage for one participant. Red dots are selected electrodes.

Participants maintained gaze position on a central fixation point by viewing a rear-projection screen using a VPixx projector (VPixx, Saint-Bruno, Québec, Canada) through a front-surface mirror attached to the head coil (screen resolution 1920 × 1080 pixels; 83.5 cm viewing distance, total screen size subtending ∼43° × 24°).

### Experimental procedure

To localize somatosensory cortex (S1) participants completed two runs of a functional finger localizer. Vibrotactile stimuli were 5-second vibrations at a 110 Hz carrier frequency starting at 0 phase, amplitude-modulated at 1 Hz by a sinusoidal envelope ranging from 0 to 1 starting at 0 phase. The stimulus was scaled to a peak amplitude of 2 V (maximum output) and delivered to the fingertip pads at half strength. Prior to scanning, we ensured that all stimulators could be felt by the participant, and the intensity of stimulation on each fingertip was reported to be similar. Tactile stimuli swept across the fingers, moving from one finger to the next every five seconds in ascending order (from thumb to little finger). The total run duration was 260 s and consisted of 10 full sweeps (thumb - little finger) with a 5 s blank at the beginning and the end. We modeled these data using a phase-encoded analysis (Engel et al., 1994; Sanchez-Panchuelo et al., 2010; Sereno et al., 1995). The goodness-of-fit (coherence) of the model to the BOLD time course was used to identify fingertip-selective voxels within somatosensory cortex (**Figure 1B**).

In the main experiment, each participant completed 8–9 runs in one scan session (lasting ∼1.5 h). On each trial, identical vibrotactile waveforms (110 Hz carrier, starting at 0 phase) were delivered simultaneously to all five fingertips of the non-dominant hand, with three stimulators per fingertip (15 stimulators total). Unlike the localizer, in which stimulation swept across fingers and the 110 Hz carrier was amplitude-modulated at 1 Hz by a sinusoidal envelope, pulses in the main experiment were delivered simultaneously to all fingertips and defined by a rectangular envelope that simply turned the carrier amplitude on and off. The carrier was scaled to a peak amplitude of 2 V (maximum output) and delivered to the fingertip pads at half strength. Prior to scanning, we ensured that all stimulators could be felt by the participant, and the intensity of stimulation on each fingertip was reported to be similar. The experiment used vibrotactile stimuli that can be grouped into two categories (**Figure 2A**): 1) 7 single-pulse vibrotactile stimuli varying in duration (0, 0.05, 0.1, 0.2, 0.4, 0.8 or 1.2 s); 2) 7 paired-pulses in which two vibrotactile stimuli of a fixed duration (0.2 s each) were separated by a varying inter-stimulus interval (0, 0.05, 0.1, 0.2, 0.4, 0.8 or 1.2 s). Note that this makes the 0.4 s single-pulse and the 0 s inter-stimulus interval paired-pulse stimulation identical, leading to a total of 13 unique tactile stimuli.

**Figure 2.**
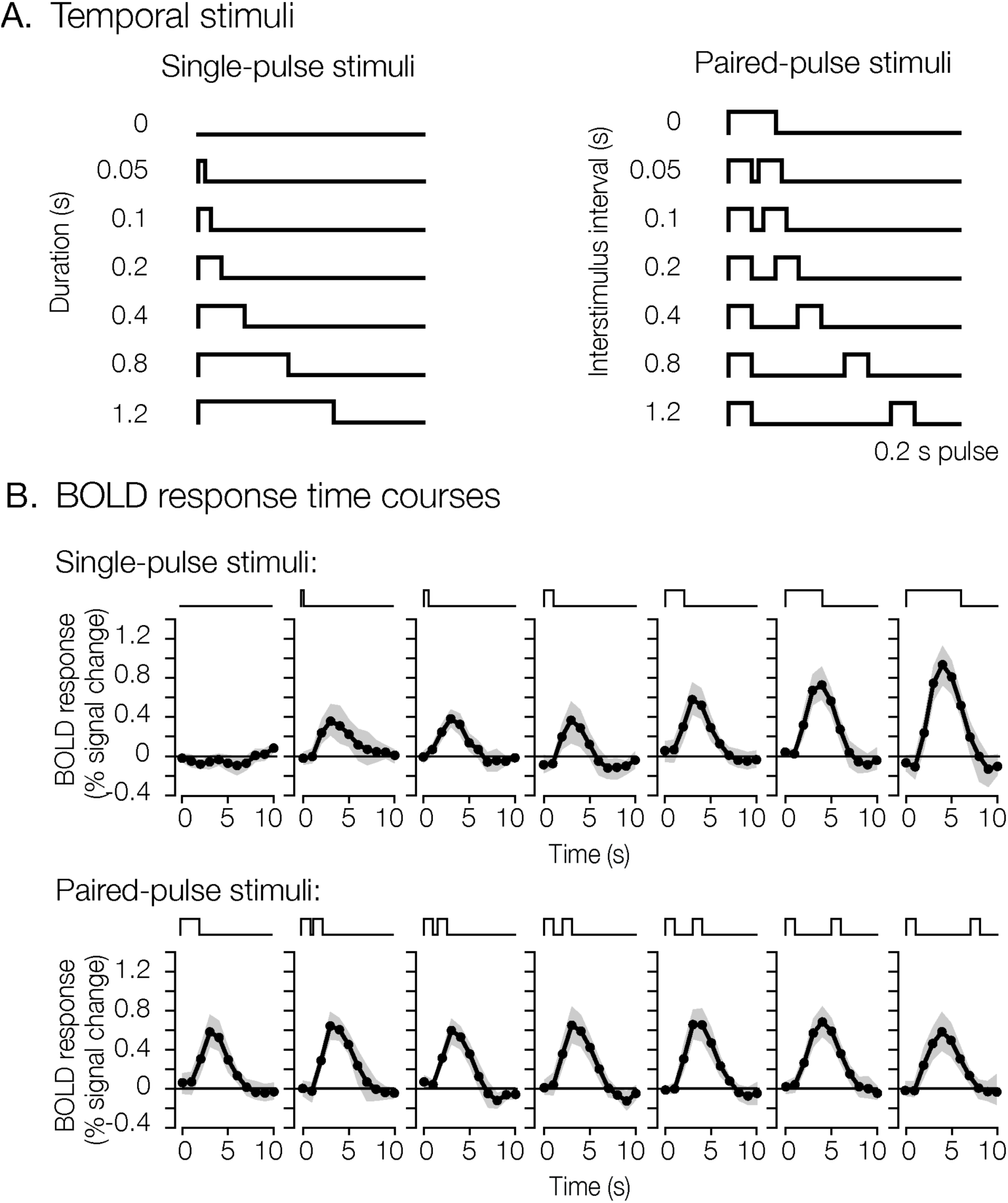
Temporal stimuli and measured BOLD response time courses in somatosensory cortex. **A**. Vibrotactile stimuli were presented at 110 Hz. The stimuli are organized into 2 categories: single-pulse stimuli, which vary in total duration and paired-pulse stimuli of 0.2 s each where the inter-stimulus interval varied. In the figure, the different stimuli are shown schematically as square-wave profiles. **B**. Group BOLD response time courses. Shaded area indicates the bootstrapped 95% confidence interval. This figure and all following figures can be reproduced with the code contained in the createFigures folder in https://github.com/ilonabloem/tactileTemporalNormalization. This figure is produced with show_figure2.m.

We used an event-related design with a variable inter-trial interval (ranging between 3–15 s). We optimized the event order and blank durations using FreeSurfer’s Optseq2 scheduling tool (Dale, 1999). To ensure a stable baseline, we additionally added a 10 s blank period to the start and end of each run. The total run duration was 320 s and consisted of 39 events (3 repetitions of each unique stimulus type). Throughout an experimental run, participants performed a fixation-color change-detection task to keep their attentional state stable. The color of the central fixation point (0.15 degree visual angle) changed between white and red (ranging between 15–29 changes per run). The time between subsequent color changes was based on an exponential distribution (mean: 10 s), offset to enforce a minimum interval of 3 s. Change times were rounded to the display’s refresh rate, and any changes that would occur in the last 3 s of a run were omitted. This procedure ensured the onset of the fixation color changes was independent of the tactile stimulus onsets.

To minimize the potential confound of hand movements related to the fixation task, participants were instructed to count how often the fixation color changed throughout a run without making responses during the run. At the end of the run, participants viewed a four-choice display and they selected an answer by using a MR-safe button box. Participants were able to do this task as reflected by the percentage of correct responses being far above chance (average across participants: 41.7%, chance level is 25%), and a small difference between the reported and actual number of color changes (mean absolute error ranged between 0.6 and 2.2 changes across participants).

### MRI data acquisition

All MRI data were collected using a Siemens MAGNETOM Prisma 3 Tesla scanner (Siemens, Erlangen, Germany) with a 64-channel head coil at New York University’s Center for Brain Imaging. fMRI data were acquired with a simultaneous multislice (SMS) gradient echo-planar acquisition protocol (Moeller et al., 2010; Xu et al., 2013): 2 mm isotropic voxels; FOV = 208 x 208 mm, 66 slices; TR = 1000 ms, TE = 37.60 ms, flip angle = 68°; multiband acceleration factor 6. We acquired two spin echo echo-planar scans with opposite y-axis phase encoding directions to allow for susceptibility distortion correction (2 mm isotropic voxels; FOV = 208 x 208 mm, 66 slices; TR = 6350 ms, TE = 51.2 ms, flip angle = 90°). During a separate scan session, we acquired a whole-brain anatomical scan using a T1-weighted 3D MPRAGE sequence: 0.8 mm isotropic voxels; FOV = 256 x 240 mm; 208 slices; TR = 2400 ms; TE = 2.24 ms; TI = 1060 ms; flip angle = 8°.

### Preprocessing using fMRIPrep

fMRI data were preprocessed using fMRIPrep 23.1.2 (Esteban et al., 2019), which is based on Nipype 1.8.6 (Gorgolewski et al., 2011).

#### Anatomical data preprocessing

The T1-weighted (T1w) image was corrected for intensity non-uniformity with N4BiasFieldCorrection (Tustison et al., 2010), and skull-stripped. The anatomical image was segmented into cerebrospinal fluid, white matter and gray matter using FSL *fast* (Y. Zhang et al., 2001). Cortical surfaces were reconstructed using recon-all in FreeSurfer 7.3.2 (Dale et al., 1999).

### Functional data preprocessing

For all functional runs, we applied the following preprocessing steps. First, a reference volume and its skull-stripped version were generated. Head-motion parameters with respect to the BOLD reference (transformation matrices, and six corresponding rotation and translation parameters) were estimated before any spatiotemporal filtering using *mcflirt* (FSL; Jenkinson et al., 2002). A B0-nonuniformity map was computed from the AP and PA phase encoding fieldmap scans with *topup* (FSL; Andersson et al., 2003), and aligned to the reference volume with rigid registration. BOLD runs were slice-time corrected and co-registered to the T1w reference using boundary-based registration (FreeSurfer; Greve & Fischl, 2009). The resulting transforms, together with motion and distortion correction, were then applied in a single resampling step to bring the functional data into T1w anatomical space.

### fMRI data analysis

We modeled the BOLD time course of each tactile stimulus condition by using a finite-impulse response (FIR) modeling approach (Dale, 1999; Glover, 1999) implemented in *GLMdenoise* (window length 11 s; Kay et al., 2013), applied to all voxels within the cortical grey matter ribbon. Noise regressors were derived by applying PCA to the time-series of the cortical ribbon voxels whose responses were unrelated to the experimental paradigm; the optimal number of principal components was determined automatically via cross-validation (Kay et al., 2013). This deconvolution analysis provided 11 beta weights describing the time course of the BOLD response for each tactile stimulus condition for each voxel.

### Voxel Selection

We used the Glasser atlas (Glasser et al., 2016) in combination with the tactile fingertip localizer experiment to select voxels within the somatosensory cortex. The primary somatosensory cortex encompasses regions BA3b, BA1, and BA2 from the Glasser atlas (Glasser et al., 2016; Kaas, 1993; Schellekens et al., 2021). Within this larger somatosensory region of interest (ROI) voxel selection was restricted to voxels with a coherence of 0.25 or greater from the phase-encoded localizer analysis. These selected voxels were averaged to obtain a participant-level BOLD time course for each tactile stimulus condition. Group-level BOLD response time courses were computed by averaging across participants and variability was estimated via bootstrapping over participants independently for each condition (500 resamples). From the bootstrapped data we computed confidence intervals. The same set of bootstrapped group-average time courses was used as input for the model fits, allowing us to obtain corresponding bootstrap distributions over model parameters.

## 3. iEEG Methods

### Participants

Intracranial EEG (iEEG) data were measured from two participants undergoing subdural electrode implantation for clinical purposes, one at the University Medical Center Utrecht (UMCU) and the other at New York University (NYU) Grossman School of Medicine. Written informed consent to participate in this study was given by both patients. The study was approved by the NYU Grossman School of Medicine Institutional Review Board and the ethical committee of the UMCU, in accordance with the Declaration of Helsinki (2013). The participant at NYU was implanted with standard clinical sEEG depth electrodes, while the participant at UMCU was implanted with clinical subdural grid electrodes (**Figure 1c**). Electrode implantation and locations were guided solely by clinical requirements.

### Experiment procedures

The vibrotactile stimuli, apparatus, experimental design, and procedures were the same as the fMRI experiment, except that the UMCU participant was tested with only one vibrotactile stimulator per fingertip instead of three. Stimuli were generated on an Acer laptop (N16P5, 14-inch, 920×1080 resolution) running windows at NYU and on an Acer Aspire 7 at UMCU. The participant at NYU completed 5 runs of the task, which included 144 trials (12 trials per condition), taking about 45 minutes to complete. The UMCU participant completed one run of the task, which included 72 trials (6 trials per condition) and took approximately 5 minutes. Stimulus onsets were synchronized with the iEEG recordings via a serial-port connection between the laptop and the iEEG amplifier. Neural signals were acquired with a Micromed amplifier at 2048 Hz, using a 0.15 Hz high-pass filter and a 500 Hz low-pass filter.

### Data preprocessing

Data were read into MATLAB, release 2024b, using the FieldTrip Toolbox (Oostenveld et al., 2011) and preprocessed with custom scripts (https://github.com/WinawerLab/ECoG_utils). Data were separated into individual runs and formatted to conform to the ieeg-BIDS (Brain Imaging Data Structure) format (Holdgraf et al., 2019). Data were then re-referenced to the common average by subtracting the mean of all electrodes excluding those identified by the neurologists as pathological (bidsEcogRereference.m). Next, a time-varying broadband estimate was computed in the following way (bidsEcogBroadband.m). First, the voltage traces were bandpass filtered using a Butterworth filter (passband ripples, 3 dB, stopband attenuation 60 dB) for non-overlapping 10-Hz-wide bands spanning 50 to 200 Hz. Bands within 10 Hz of the main electrical line frequency or its harmonics were excluded (NYU: 70-80 Hz, 90-100 Hz, 100-110 Hz, 130-140 Hz, 140-150 Hz, 150-160 Hz, 160-170 Hz, 190-200 Hz; UMCU: 60-70 Hz, 70-80 Hz, 80-90 Hz, 110-120 Hz, 120-130 Hz, 130-140 Hz, 160-180 Hz, 180-190 Hz, 190-200 Hz). The power envelope of each bandpass-filtered voltage time course was then calculated as the square of its Hilbert transform. The broadband time series was then computed as the geometric mean of the envelopes. Because the low-pass filtering properties of neurons (e.g., dendritic leaky integration) cause field potential power to decline with frequency (Miller et al., 2014), taking the geometric mean ensures that each frequency band is treated as equally informative about the underlying neural generator.

### Data and electrode selection

The preprocessed data were analyzed using custom scripts. For each participant, we segmented the broadband time courses into epochs from −0.4 to 2.2 s relative to stimulus onset and converted them to percent signal change by dividing by and subtracting the mean prestimulus baseline. Epochs were selected for further analysis if they passed an automated artifact rejection procedure adapted from (Groen et al., 2022). This procedure was applied independently for each electrode. Epochs were included if the maximum broadband amplitude outside the stimulus presentation window (0 – 0.4 s after stimulus onset) did not exceed the average of the maximum response plus 3 SDs inside the stimulus presentation window. This removed artifacts and broadband bursts unrelated to stimulus processing. This process rejected only a small percentage of epochs: 2.8% for the NYU participant and 2.1% for the UMCU participant.

Electrode selection proceeded in two stages. First, we selected electrodes based on their evoked broadband responses, a method adapted from Zhou et al. (2019). For each electrode, we averaged epochs across stimulus conditions to obtain a single time course per electrode and focused on the response from 0 to 0.4 s relative to stimulus onset. Within this window, we computed the maximum response, and used a data-driven threshold as the median of these maxima across electrodes plus 1 SD. We excluded electrodes that fell below this threshold. Second, we inspected the anatomical locations of the remaining electrodes. Depth electrodes from the NYU participant were localized based on pre-implantation (MRI) and post-implantation (CT) structural images. Electrodes from the UMCU participant were localized from a postoperative CT scan and coregistered to the preoperative MRI (Branco et al., 2018). The selected electrodes fell within presumptive somatosensory cortex: the postcentral gyrus or supramarginal gyrus in the Freesurfer cortical parcellation (Fischl, 2004). In total this procedure resulted in 14 electrodes for the NYU participant and 5 electrodes for the UMCU participant (**Figure 1c**).

### Average responses and bootstrapping

We computed a group average broadband time course for each stimulus condition using a bootstrap procedure. For each participant, we first performed 1000 bootstrap resamples by drawing *N* electrodes with replacement from the *N* available electrodes for each participant (*N* is 14 for the NYU participant and *N* is 4 for the Utrecht participant). We then averaged the broadband time series across the selected electrodes separately for each participant, yielding 1000 time courses per stimulus condition and participant. For each bootstrap iteration, we then averaged the time courses across participants, so that the two participants’ data were weighted equally. This bootstrap distribution of group-average responses was subsequently used to derive confidence intervals on the mean broadband power time courses and to estimate variability in the fitted model parameters across bootstraps.

## 4. Temporal dynamics models

We modeled the data from both the fMRI and the iEEG experiments with a time-invariant linear model (i.e., convolution with a linear filter) and with a nonlinear (normalization) model. The models take the envelope of the vibrotactile stimulation as input and predict either the BOLD time course or the iEEG broadband time course as output.

### Linear model

In the linear model, the predicted neural response (*R_linear_*) was computed by convolving the stimulus time courses (*S*) with a temporal impulse response function (*k*):

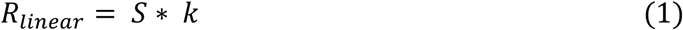

The physical stimulation consisted of a continuous 110 Hz vibration whose amplitude was gated on and off by a rectangular envelope defining the onset and offset times of each pulse. In our models, the stimulus time course is represented by this envelope as a binary vector that takes the value 1 during vibrotactile stimulation and 0 otherwise, treating each pulse as a square wave in time. For fMRI data, the BOLD measurement depends on both the neural response and the neurovascular coupling. For the linear model, we assume that both are convolutional, and so *k* incorporates both a neural impulse response function and a hemodynamic impulse response function (HIRF). Because the neural impulse response function is assumed to be brief (tens to hundreds of ms) and the HIRF long (seconds to tens of seconds), the specific shape of the neural impulse response function is unimportant for the linear model, and *k* is dominated by the hemodynamic response. We parameterized *k* as the weighted difference of two gamma functions (MATLAB function ‘gampdf’):

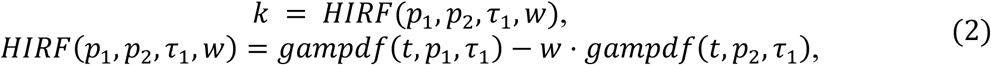

where *t* is time, *p*_1_ and *p*_2_ are shape factors affecting the positive and negative components of the temporal profile, respectively, *τ*_1_ is a scale factor fixed at 1, and *w* is the relative weight of the negative component. To match the temporal resolution of the measured fMRI time courses, *R_linear_* was downsampled by averaging within each 1-s fMRI volume.

For iEEG data, *k* was an impulse response function (IRF), implemented as a weighted difference between two gamma functions:

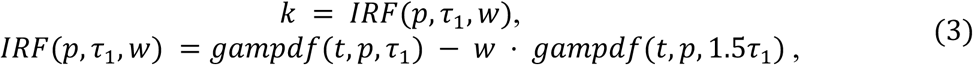

where *p* is the shape factor fixed at 2, *τ_1_* is a scale factor determining the time-to-peak, and *w* is the relative weight of the negative component.

While both the HIRF for modeling the fMRI response and the neural IRF for modeling the iEEG data were parameterized as the difference of two gamma functions, they differ in time scale by an order of magnitude: the time to peak is several seconds for the HIRF and tens to hundreds of ms for the neural IRF. Moreover, they differ in which parameters were allowed to contribute to the shape (HIRF: the shape parameter; IRF: the scale parameter). Fitting the model to the fMRI data with both *p* and *τ*_1_ as free parameters did not improve the fits.

Finally for both imaging modalities, predicted time courses were scaled by a common gain (*g*) parameter across stimulus conditions:

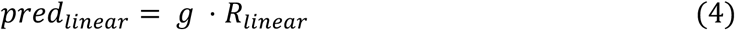

### Normalization model

We adopt two variants of the normalization model that have been previously implemented to model temporal dynamics in visual fMRI experiments, “Compressive Temporal Summation” (Zhou et al., 2018) and “Delayed normalization” (Groen et al., 2022). Both implementations differ from the linear model in that the neural response is divisively normalized. To compute the model outputs, the stimulus time course was first convolved with an *IRF* to yield a linear neural response:

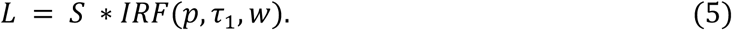

The IRF was modeled as a difference of two gamma functions (equation 3). For fMRI data, *p* was fixed to 2, and the weight for the second negative gamma function in the IRF was fixed to *w*=0, as in Zhou et al. (2018). The BOLD response was modeled as the normalized neural response convolved with the HIRF (equation 2) as follows:

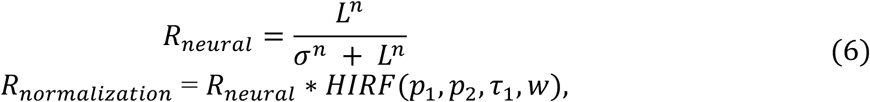

where *p*_1_, *p*_2_, *w*, and *σ* were free parameters, *τ*_1_ was fixed at 1, and the exponent was fixed at *n*=2. The normalization pool (the denominator of the divisive normalization in equation 6) consists of the response itself and a constant *σ*. When *σ* approaches 0, normalization is strong, such that the neural response rises steeply and reaches a plateau with increasing *S*. When *σ* is large, the neural response is nearly linear as a function of *S*. After convolving with the HIRF, the modeled response was downsampled by averaging within each 1-s fMRI volume.

For the iEEG data, we modeled divisive normalization with a delay as:

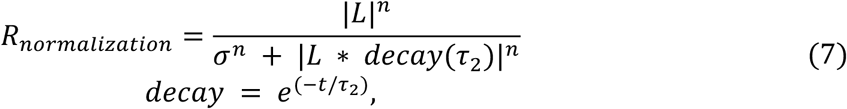

where the neural response was divisively normalized by a low-pass filtered version of itself. This was implemented as a convolution with an exponential decay function with time constant *τ_2_*. The decay term reflects the observation that the normalization response is slower than the linear response, a phenomenon that is evident in the iEEG signal (Zhou et al., 2019) but not in the slower fMRI signal.

Finally for each measurement modality, the predicted time course was scaled by a gain factor (*g*):

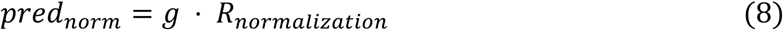

### Model fitting and parameter estimation

We fit models separately to the time course of the BOLD responses and the time course of the iEEG broadband responses. For each, we minimized the root-mean-squared error (RMSE) between model predictions and the data, concatenated across temporal conditions, using the Bayesian Adaptive Direct Search (BADS; Acerbi & Ma, 2017) algorithm. For the fMRI data, the linear model has a total of 4 free parameters (*p_1_*, *p_2_*, *w*, *g*), while the normalization model has 6 free parameters (*σ, τ_1_*, *p_1_*, *p_2_*, *w*, *g*). For the iEEG data, the linear model has a total of 3 free parameters (*τ_1_*, *w*, *g*), while the delayed normalization model has 6 free parameters (*σ, τ_1_*, *w*, *τ_2_*, *n*, *g*). To compare models while minimizing the effect of unequal numbers of free parameters, we quantify model accuracy by cross-validated *R*^2^ (the explained variance) of model predictions using a leave-one-condition-out approach.

### Data availability

All code used for the purpose of this article can be found at https://github.com/ilonabloem/tactileTemporalNormalization. Both the fMRI and iEEG processed datasets are available on the Open Science Framework (https://osf.io/f6tzr).

## 5. Results

We measured BOLD responses in the somatosensory cortex while vibrotactile stimulation (110 Hz) was applied to all five fingertip pads of the participants’ nondominant hand. We presented seven single-pulse trial types and seven paired-pulse trial types (**Figure 2A** and Methods). We used an event-related design with a variable inter-trial interval (3 to 15 s) while participants were engaged in a temporally independent fixation dot-color change-detection task. The BOLD response time course of each vibrotactile stimulus was estimated using a finite impulse-response (FIR) modeling approach (Kay et al., 2013). This provided, for each tactile stimulus condition, a time course composed of 11 beta-weights starting at stimulus onset (**Figure 2B**).

### Temporal summation in somatosensory cortex is sub-additive

The temporal properties of the BOLD response are often assumed to be linearly related to stimulus duration. Seminal studies (Boynton et al., 1996, 2012) showed that BOLD responses to longer stimuli can be accurately predicted from shorter duration stimuli. However, recent work in visual cortex revealed that at shorter timescales, BOLD responses exhibit complex temporal dynamics (Birn & Bandettini, 2005; Kupers et al., 2024; Lewis et al., 2018; Siero et al., 2013; Stigliani et al., 2019; Yeşilyurt et al., 2008; Zhou et al., 2018). Specifically, studies report sub-additive temporal summation, where responses to combined stimuli are less than the sum of individual responses. To investigate whether BOLD responses in somatosensory cortex exhibit similar sub-additive temporal effects, we simulated predicted BOLD time courses assuming a linear shift-invariant system. For such a system, the response evoked by a 0.4 s stimulus should be identical to the sum of two 0.2 s stimulus responses, with the second stimulus shifted by 0.2 s. However, if responses are nonlinear or sub-additive, this linear prediction will overestimate the measured responses. **Figure 3** illustrates that the predictions of a linear shift-invariant system consistently overestimate the measured responses to the single-pulse stimuli. The discrepancy is largest when predicting a long-duration stimulus from a short-duration stimulus, as illustrated by the bottom left panel where a 0.8 s stimulus is poorly predicted by shifted and summed copies of the 0.05 s stimulus. However, this discrepancy is present across all duration combinations. This demonstrates that temporal summation in somatosensory cortex is sub-additive: doubling the duration of the stimulus does not result in a doubling of its response.

**Figure 3.**
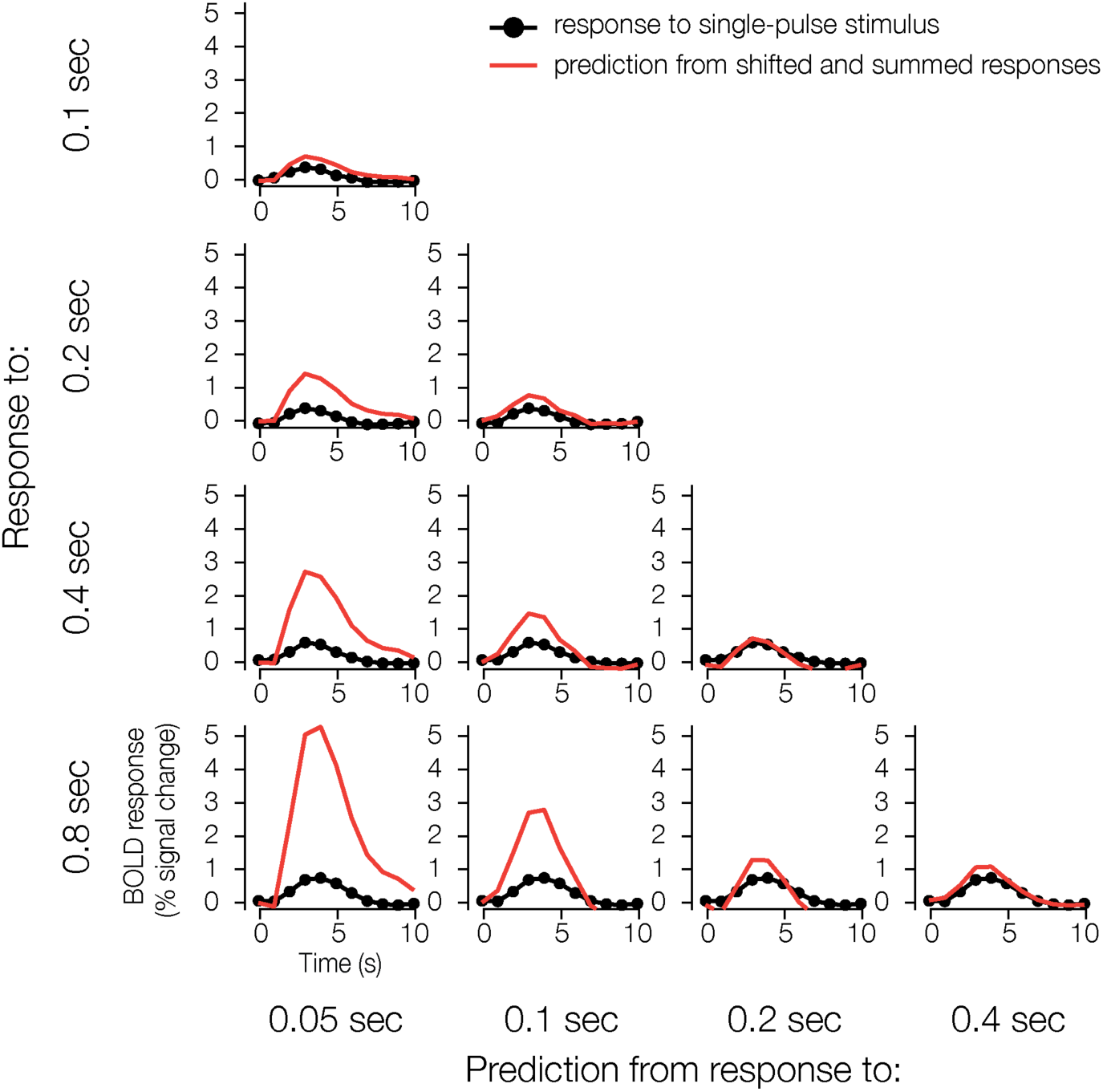
Sub-additive temporal summation measured with fMRI. Somatosensory cortical responses (black) are much less than the predictions from shifted and summed responses of shorter stimuli (red). Figure conventions adapted from Boynton et al. (1996). This figure is produced with show_figure3.m.

### Temporal subadditivity is captured by a normalization model

To quantify the temporal dynamics of the somatosensory BOLD response time courses, we compared the accuracy of model predictions between a linear model and a normalization model. Variants of these models have been previously used to assess temporal dynamics within visual cortex (Chapman & Denison, 2025; Groen et al., 2022; Lisberger & Movshon, 1999; Zhou et al., 2019). For our implementation, both models start with the time course of the vibrotactile stimulus (at millisecond resolution) as input and produce a predicted response time course (at second resolution) as output (**Figure 4**).

**Figure 4.**
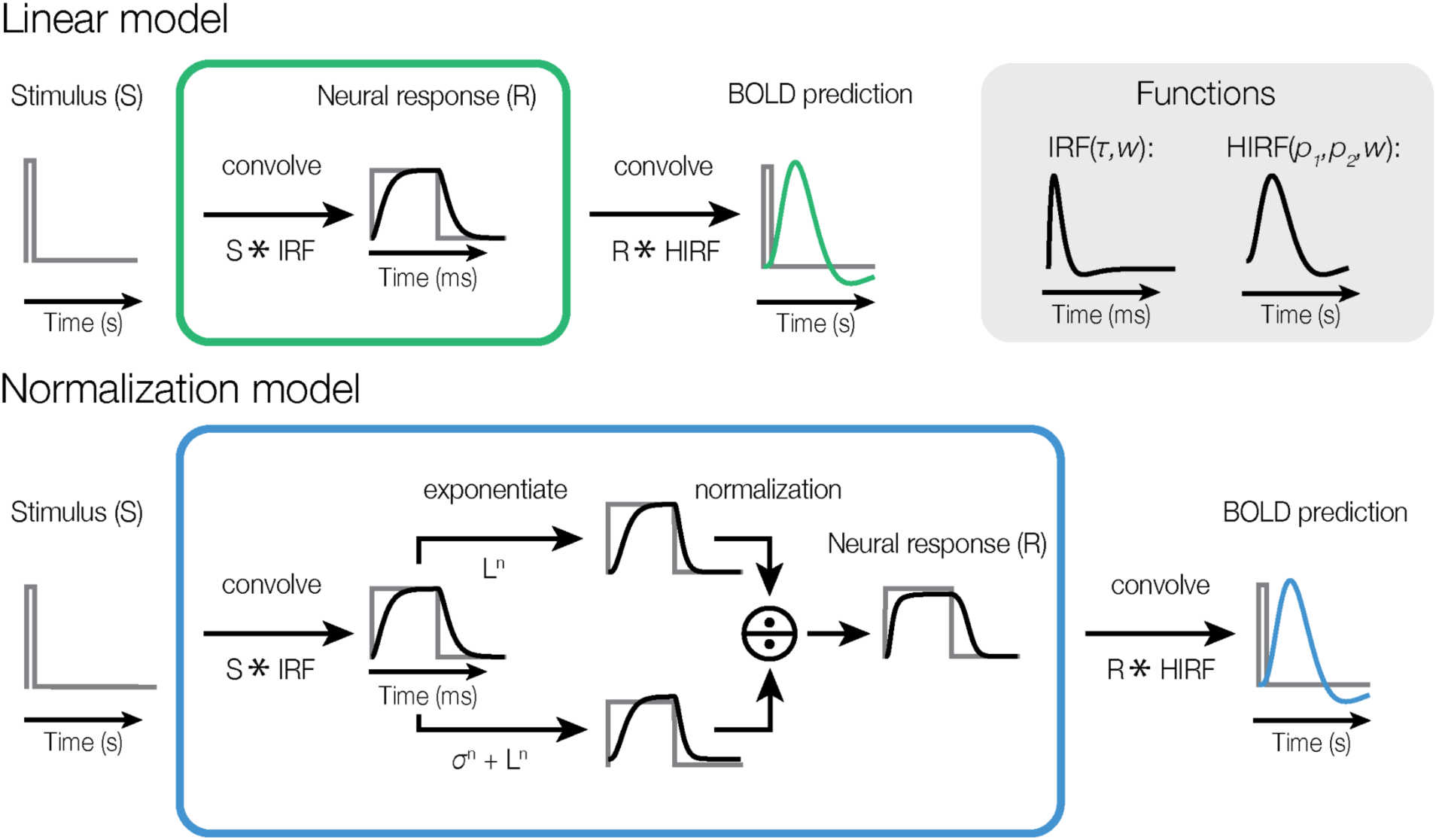
Temporal dynamics models. In the linear model (top left panel), the stimulus time course S is convolved with a neural IRF (green box), and then with the HIRF. The HIRF is outside the green box to indicate that it is hemodynamic and not neural. The top right panel shows the two convolution kernels, the neural IRF and the HIRF. The neural IRF is an order of magnitude shorter. Note that the neural convolution stage (green box) is shown for completeness but was omitted from the fitting procedure because two successive convolutions are equivalent to a single convolution. In the normalization model (bottom panel), the stimulus time course is convolved with a neural IRF, followed by divisive normalization, followed by convolution with an HIRF. As with the upper panel, the blue box represents the neural model. This figure is produced with show_figure4.m.

In the linear model, the stimulus time course was transformed to a predicted BOLD response by convolving the stimulus time course with a hemodynamic response function (HIRF). Because any linear temporal filtering at the neural level is captured by the effective HIRF, we did not include a separate neural impulse response. In contrast, the normalization model first convolved the stimulus time course with a neural IRF, after which the neural response undergoes divisive normalization (Carandini & Heeger, 2012). This involved dividing the exponentiated neural response by a normalization pool consisting of itself plus an exponentiated semi-saturation constant. The output of this normalization stage was subsequently convolved with the

HIRF and downsampled to generate a predicted BOLD time course at the 1-s resolution of the fMRI measurement. Each model was fit to the BOLD response time courses for all tactile stimulus conditions simultaneously.

To assess model accuracy, we computed cross-validated variance explained (*R^2^_crossval_*) by a leave-one-condition-out approach. The normalization model provided more accurate predictions of the BOLD responses elicited by all vibrotactile stimuli compared to the simpler linear model (**Figure 5A**). Across all tactile stimuli, the normalization model reached a 0.91 *R^2^_crossval_*, whereas the linear model reached 0.67. As demonstrated in **Figure 3**, a linear prediction extrapolated from a short stimulus would inherently overestimate the response to longer or paired-pulse stimuli due to sub-additivity. However, because we fit the linear model to all single- and paired-pulse conditions simultaneously rather than extrapolating from a short stimulus, the model is forced to find a compromise. To minimize large errors from overpredicting the longest single-pulse stimuli, the model reduces its overall gain. Consequently, this compromised fit tends to underestimate the responses to paired-pulse stimuli and shorter single-pulse stimuli (< 0.4 s), while overestimating the longer single-pulse conditions. In contrast, the normalization model does not require this compromise. Because the neural response is divided by itself (plus a semi-saturation constant), the model inherently compresses the output for larger inputs. **Figure 5B** shows the bootstrapped parameter estimates for both models. Notably, the semi-saturation constant (*σ)* estimated by the normalization model is close to zero, indicating strong normalization.

**Figure 5.**
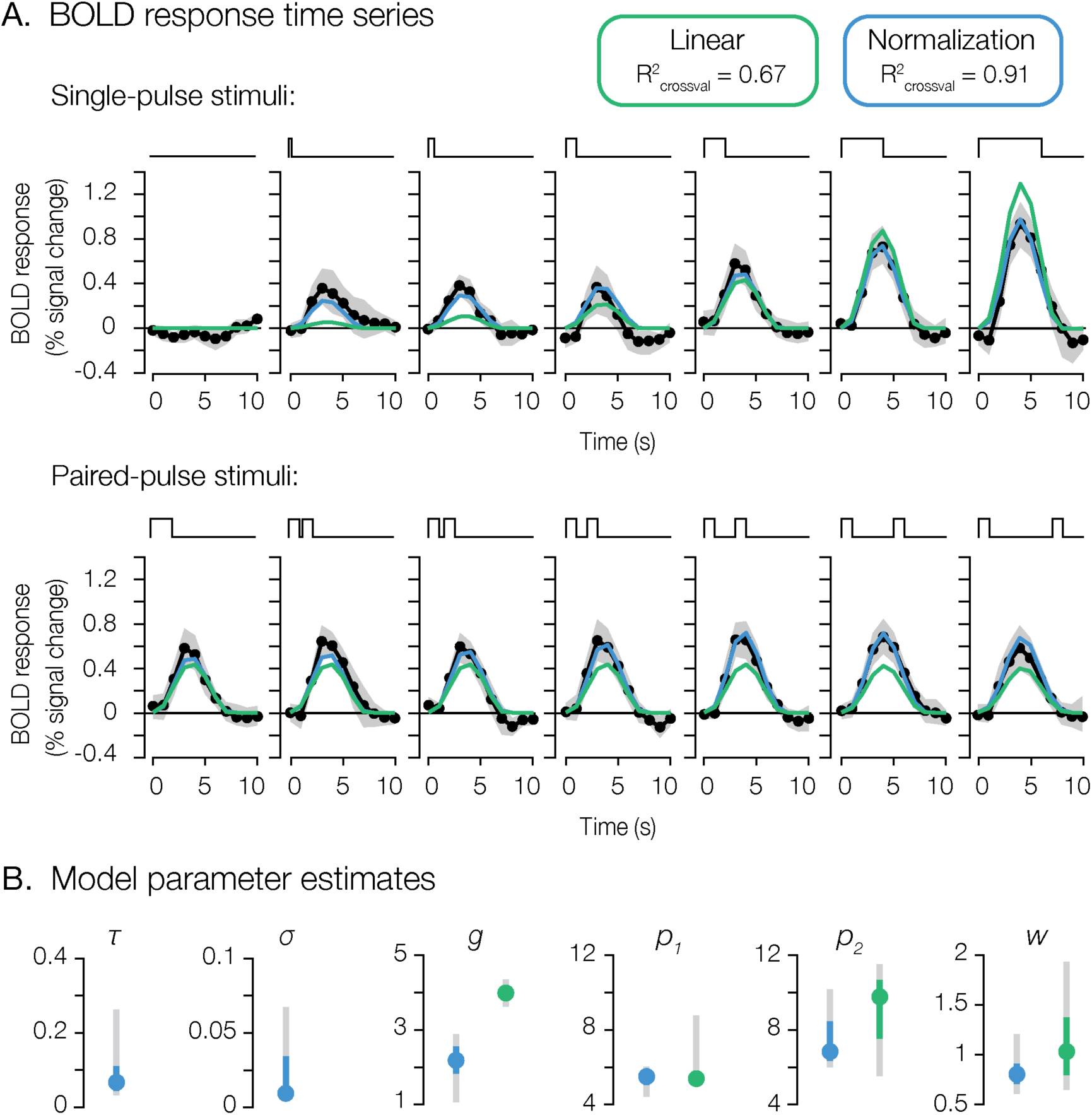
**A**. Group BOLD response time series and model predictions. Black dots are the group-averaged data, and gray shaded area is the bootstrapped 95% confidence interval. The green line represents the linear-model fit, and the blue line represents the normalization-model fit. Durations for the upper row, and inter-stimulus intervals for the bottom row, from left to right are 0, 0.05, 0.1, 0.2, 0.4, 0.8 or 1.2 s. **B**. Bootstrapped parameter estimates and confidence intervals for key model parameters: *τ* (IRF time constant), *σ* (normalization semi-saturation constant), *g* (overall gain that scales the response to all stimuli), and *p_1_*, *p_2_*, *w* (defining the HIRF). The linear model is indicated in green, and the normalization model in blue. Error bars are bootstrapped 68% (color) and 95% (gray) confidence intervals. This figure is produced with show_figure5.m.

### Summed BOLD responses reveal temporal sub-additivity

Next, we computed a summed response for both the BOLD response and model prediction time series (**Figure 6**). The use of this summed metric offers an alternative manner in which to interpret the results presented in **Figure 5**, by collapsing the time series into a single measure of total response for each condition. The linear model overpredicts responses of one-pulse stimuli longer than 0.4 s and underpredicts all two-pulse responses. The normalization model better captures sub-additive temporal summation in summed responses as stimulus duration increases, as well as the subtle increase in summed response as the inter-stimulus interval increases.

**Figure 6.**
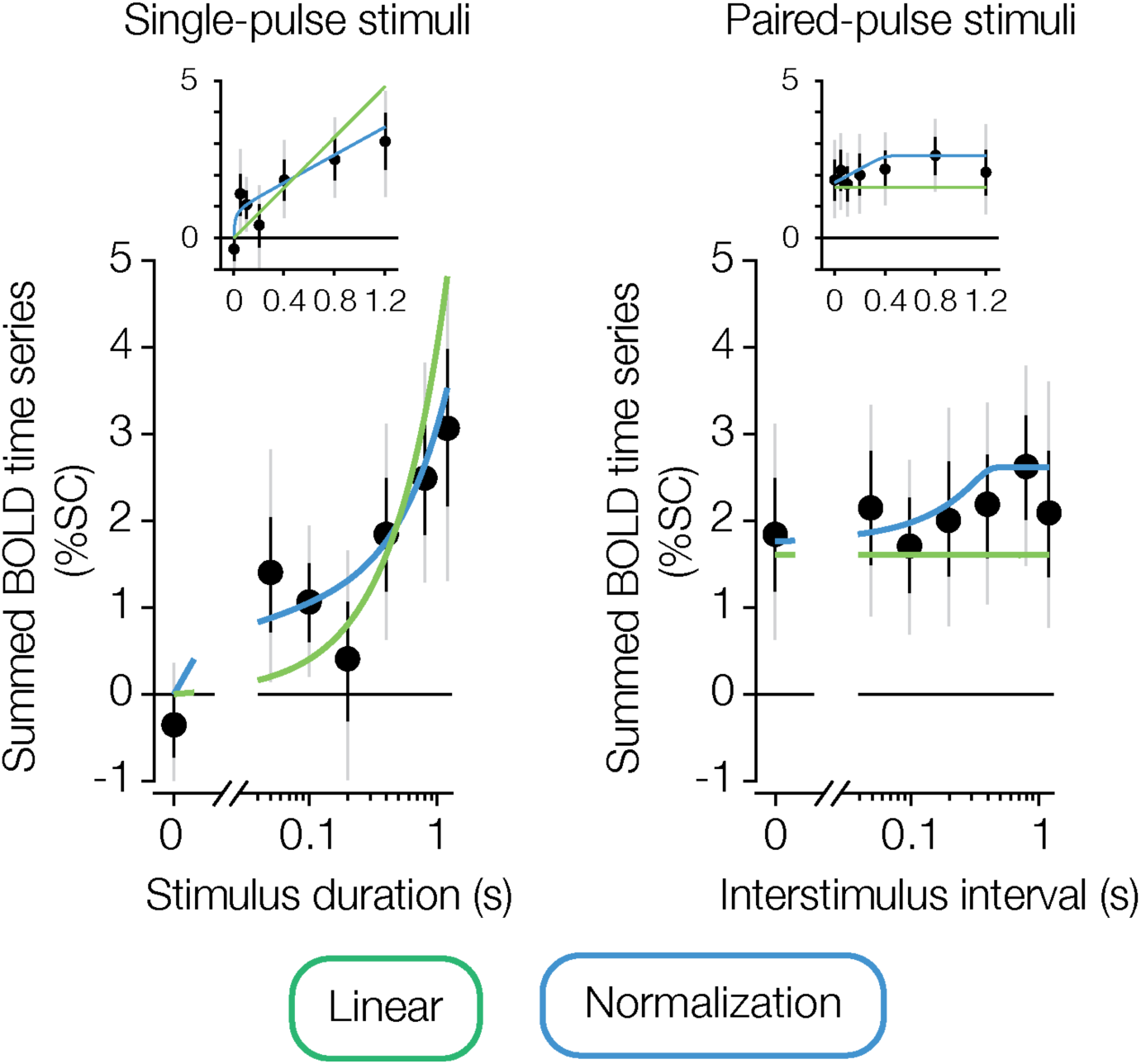
Summary metric of BOLD response and model predictions. We summed the BOLD response across the time course of each stimulus condition. These responses are visualized as a function of duration (left panels) and interstimulus interval (right panels). Black dots represent group-averaged summed responses, with error bars indicating the bootstrapped 68% (black) and 95% (gray) confidence intervals. The lines are the summed model predictions for the linear model (green) and the normalization model (blue). Insets show the same data but on a linear *x*-axis. This figure is produced with show_figure6.m.

### iEEG responses in somatosensory cortex confirm sub-additive temporal dynamics

To assess temporal dynamics at a much higher temporal resolution, and to determine whether the observed temporal subadditivity in the fMRI experiment reflects nonlinearities in the neural responses rather than in the hemodynamic response function, we turned to intracranial EEG. In particular, the BOLD response cannot distinguish whether the observed non-linearity influences the initial transient neural response, the ongoing sustained response, or a combination of both as proposed by two-channel models of temporal processing (Horiguchi et al., 2009; Stigliani et al., 2017). We recorded broadband power responses in somatosensory cortex from 2 participants (19 electrodes in total) using the same vibrotactile stimuli, apparatus, experimental design, and procedures as in the fMRI experiment, except that the interval between trials was shorter (**Figure 7A**). Broadband power is interpreted as reflecting local population cortical activity (Miller et al., 2014), making it well suited to test whether temporal sub-additivity is present at the neural level.

**Figure 7.**
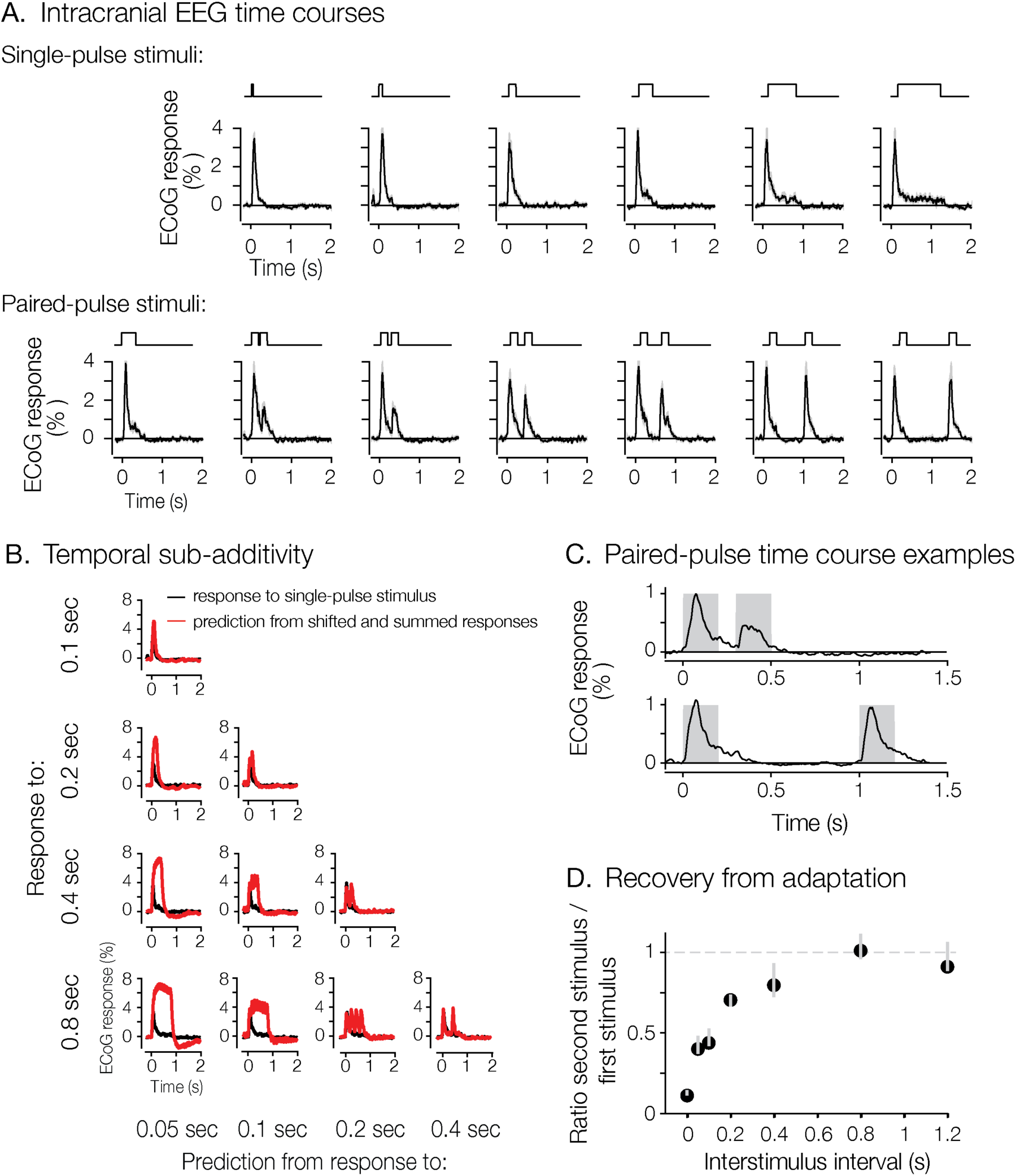
iEEG responses in somatosensory cortex are sub-additive. **A.** Average participant iEEG responses to stimuli that varied in duration and inter-stimulus interval. **B.** iEEG responses (black) are less than the predictions from responses of shorter stimuli (red) as a linear shift-invariant system. **C.** Example of adaptation to two paired-pulses of tactile stimuli with equal duration (0.2 s each) separated by either a short (0.1 s, upper) or a long (0.8 s, lower) inter-stimulus interval. The iEEG time-courses (black thick lines) show that the response to the second stimulus is reduced for a short inter-stimulus interval whereas it recovers for the longer interval. Gray shading indicates stimulus presentation. **D.** The ratio of the second pulse response to the first pulse response as a function of inter-stimulus interval. Error bar denotes the 68% CI. This figure is produced with show_figure7.m.

The iEEG time courses revealed temporal dynamics consistent with those measured by fMRI. First, when stimulus duration was doubled, the neural response did not double as would be predicted from a linear system (**Figure 7A**). This sub-additivity is further illustrated in **Figure 7B**: neural responses consistently fell far below the predictions of a linear, shift-invariant system, which were generated by shifting and summing responses from shorter durations. Second, iEEG made it possible to observe recovery from adaptation at the millisecond scale. For instance, for paired-pulse stimuli the response to the second stimulus was reduced when the inter-stimulus interval was 0.1 s but recovered to the same amplitude as the first pulse when the interval increased to 0.8 s (**Figure 7C**). **Figure 7D** quantifies this effect across all paired-pulse conditions. The ratio of the response to the second stimulus relative to the first stimulus increased from 0.2 at short inter-stimulus intervals and plateaued around 1 for intervals exceeding 0.8 s. This indicates that the response to the second stimulus at a longer inter-stimulus interval is no longer influenced by the first. This strong ISI-dependent suppression of the second pulse confirms that the temporal non-linearities are driven, at least in part, by adaptation of the transient neural response.

To model the temporal dynamics of the iEEG responses, we again compared a linear and a normalization model (see Temporal dynamics models). For iEEG time courses, the linear model convolved the stimulus time course with a neural IRF (difference of gamma functions, parametrized by a time constant and weight that determined the contribution of the second negative gamma). The delayed normalization model resembled the normalization model used for the fMRI data but additionally implemented an exponential decay function in the normalization pool (**Figure 8A**). This made the normalization model history-dependent, enabling the model to capture both the transient-to-sustained response profile, and the influence of adaptation seen in the paired-pulse conditions.

**Figure 8.**
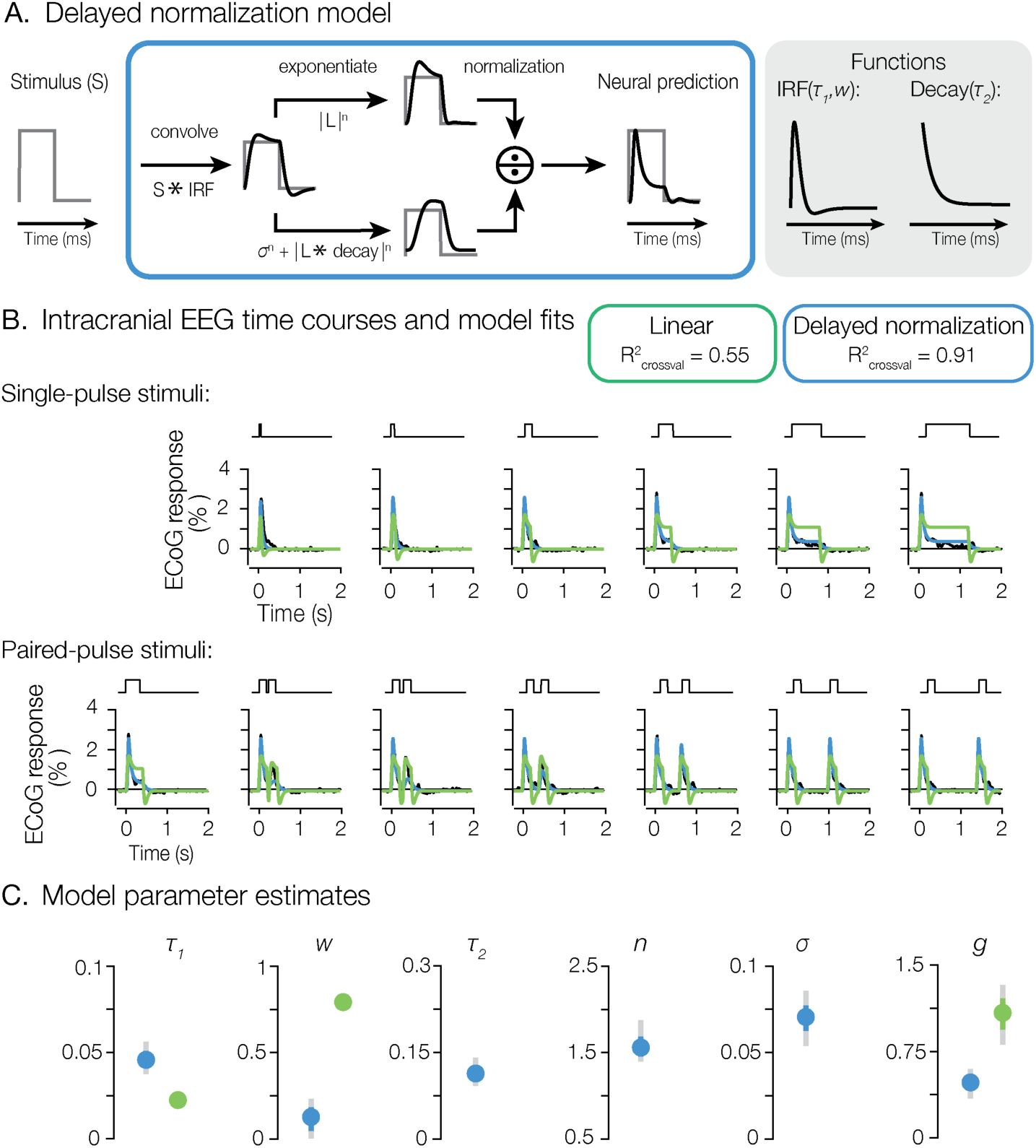
**A.** iEEG delayed normalization model. **B.** Group iEEG broadband response time series and model predictions. Black lines are the group-averaged data, the green line represents the linear-model fit, and the blue line represents the normalization-model fit. Durations for the single-pulse stimuli, and inter-stimulus intervals for paired-pulse stimuli, from left to right are 0, 0.05, 0.1, 0.2, 0.4, 0.8 or 1.2 s. **C**. Bootstrapped parameter estimates and confidence intervals for key model parameters: *τ_1_* and *w* (IRF time constant and weight), *τ_2_* (exponential-decay time constant), *σ* (normalization semi-saturation constant), *n* (normalization exponent), and *g* (overall gain that scales the responses to all stimuli). The linear model is indicated in green, and the normalization model in blue. Error bars are bootstrapped 68% (color) and 95% (gray) confidence intervals. This figure is produced with show_figure8.m.

Model predictions for the iEEG time courses closely paralleled the fMRI findings. Using leave-one-condition-out cross-validation, the delayed normalization model captured the broadband time courses to the different tactile stimuli substantially better than the linear model (**Figure 8B**; normalization model cross-validated R^2^ = 0.91, linear model: 0.55). The delayed normalization model better accounted for both the initial transient response and the subsequent adaptation in the broadband time course. While the linear model can predict an initial transient because of the biphasic IRF, it also predicts an equally large negative deflection at stimulus offset, as a single convolution cannot capture a sharp onset transient without a strong negative component. This drives the linear prediction below baseline at stimulus offset, a pattern that does not occur in the data. **Figure 8C** illustrates the bootstrapped parameter estimates for both models. Taken together with the fMRI results, this suggests that the advantage of the normalization model for both iEEG and fMRI data reflects nonlinearities in the underlying neural responses, rather than being primarily driven by properties of the hemodynamic response.

### Delayed normalization iEEG predictions explain fMRI BOLD responses

Finally, we asked whether the model predictions from the iEEG time courses could also account for the fMRI summed responses (from **Figure 6**), despite being obtained in different participants with different imaging methods. To place both measures on a common scale, we first computed a single scalar as the ratio between the summed fMRI responses and the summed iEEG responses across all conditions. This scalar represents a simple linear mapping from iEEG broadband power to BOLD response. We then multiplied the iEEG model predictions by the scalar, yielding a rescaled summed prediction that could directly be compared to the summed BOLD responses. Although the delayed normalization model was estimated based on the iEEG time courses alone, it accurately predicted the BOLD responses (*R^2^* = 0.77; **Figure 9**). The linear model fit to the iEEG data was a poor predictor of the BOLD responses (*R^2^* = 0.28).

**Figure 9.**
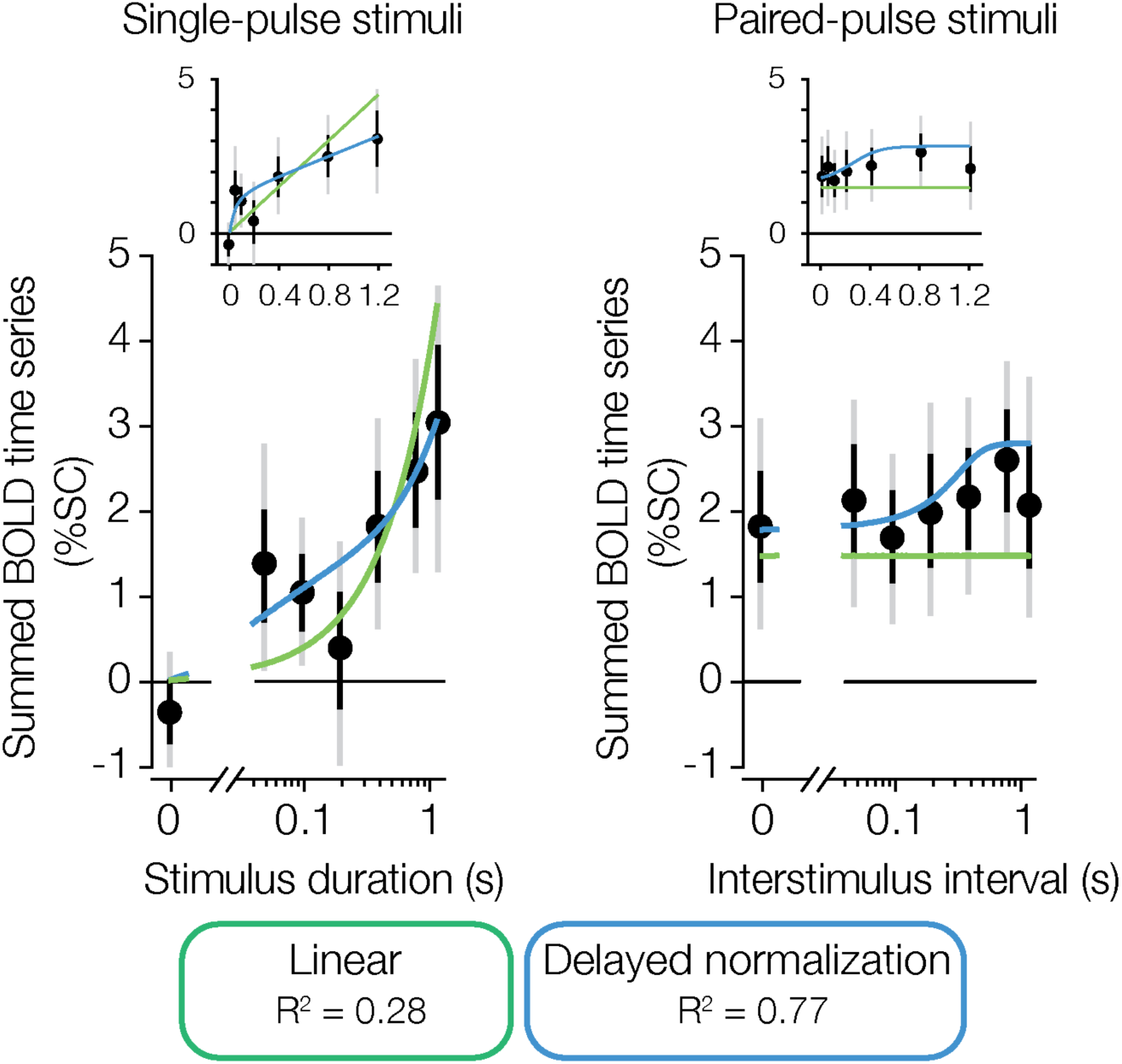
Predicting summed BOLD responses from iEEG model predictions. Summed fMRI BOLD responses (replotted from Figure 6) against a scaled and summed iEEG model prediction for the same tactile condition. Black dots represent group-averaged summed responses, with error bars indicating the bootstrapped 68% (black) and 95% (gray) confidence intervals. The lines are the scaled and summed iEEG model predictions for the linear model (green) and the delayed normalization model (blue). Insets show the same data but on a linear *x*-axis. This figure is produced with show_figure9.m.

## 6. Discussion

Neural responses in human somatosensory cortex showed robust sub-additive temporal summation for prolonged and repeated vibrotactile stimuli. Using fMRI, we found that BOLD responses scaled sub-additively with stimulus duration and repetition, indicating that temporal integration is not linear. A divisive normalization model provided a better account of the observed dynamics than a linear prediction. Intracranial EEG (iEEG) broadband power revealed the temporal dynamics at a higher temporal resolution. These neural responses confirmed sub-additive temporal summation for the same stimulus conditions. Additionally, the iEEG results demonstrated clear transient-to-sustained response profiles and inter-stimulus interval dependent recovery from adaptation for paired-pulse conditions. These dynamics were accurately captured by a delayed normalization model. Finally, we showed that the delayed normalization iEEG model predicted the summed BOLD responses well, linking neural and hemodynamic measures of temporal integration under a common normalization framework. Together, these results indicate that temporal dynamics in human somatosensory cortex are well described by divisive normalization acting over time.

### Disentangling hemodynamic and neural contributions

BOLD responses scale approximately linear for relatively long stimuli (> 3 s), such that scaled and summed copies of a 3 s stimulus can predict the response to a 6 s stimulus well (Boynton et al., 1996). In contrast, for shorter stimuli (< 2 s) BOLD responses deviate from linearity: responses to short durations are disproportionately large relative to a linear prediction from longer durations (Birn et al., 2001; Pfeuffer et al., 2003; Yeşilyurt et al., 2008).

The origin of this nonlinearity remains debated. Some studies attribute it to vascular mechanisms that generate relatively large transients in this shorter-duration regime (de Zwart et al., 2009; N. Zhang et al., 2008). More recent work instead argues that BOLD response nonlinearities for brief stimuli (< 2 s) are primarily driven by neural nonlinearities (Gaglianese et al., 2017; Siero et al., 2013; Zhou et al., 2019), because vascular dynamics (cerebral blood flow and cerebral blood volume) are proportional to one another in this regime (Polimeni & Lewis, 2021).

Our iEEG data provide a test of whether the sub-additivity we observed in the fMRI data is neural in origin. Broadband iEEG time courses exhibited a pattern of temporal sub-additivity across stimulus duration that was similar to the BOLD responses. Furthermore, a delayed-normalization model fit only to iEEG data accurately predicted the fMRI data (**Figure 9**). The close agreement between the iEEG-based normalization-model predictions and the measured BOLD responses indicates that the temporal sub-additivity we observed in the BOLD responses in this brief-duration regime reflects neural response dynamics captured by the normalization model. Together, these findings link iEEG and fMRI measurements under a common computational framework.

The computational framework links the stimulus to the cortical response, including all of the processing in between. We remain agnostic as to whether the nonlinearity arises from peripheral mechanoreceptor adaptation, normalization within somatosensory cortex, or an accumulation across multiple processing stages; disentangling these contributions would require measurements at earlier stages of the somatosensory pathway. That said, the close correspondence between the somatosensory dynamics reported here and the cortical normalization dynamics previously characterized in the visual system (Groen et al., 2022) suggests that a substantial component of the nonlinearity arises in cortex.

### Canonical computation

Divisive normalization has been proposed as a canonical neural computation—a general operation repeated across sensory systems, brain regions, and behavioral contexts (Carandini & Heeger, 2012). The normalization model we applied to the somatosensory responses was originally developed to explain temporal dynamics within visual cortex (Brands et al., 2024; Groen et al., 2022; Zhou et al., 2018). Here, we found that a normalization model accurately captured fMRI responses also in the somatosensory cortex. The delayed normalization model for iEEG extends the ‘static’ fMRI model in two ways. First, it includes an exponential decay in the normalization pool. Second, it assumes a biphasic impulse response function to approximate the neural drive. These two additions make gain control explicitly history-dependent and allow the model to capture the transient-to-sustained dynamics as well as repetition suppression present in the iEEG time courses.

The same transient and sustained neural dynamics are presumably present in the fMRI data, and we indeed observed clear temporal differences in the BOLD response, particularly for the single-pulse conditions. However, the slow hemodynamic response smooths neural activity over seconds, so millisecond scale dynamics are largely absorbed and cannot be resolved in the BOLD response. More generally, divisive normalization can be understood as a canonical computation in which neural responses are scaled relative to the pooled activity of a broader population, with different models corresponding to different assumptions about how this pool is constructed and evolves over time. In this sense, the static normalization model used for fMRI responses, and the delayed normalization model for iEEG reflect the same underlying canonical computation, adapted to the temporal resolution of either the indirect hemodynamics of fMRI versus a more direct measurement of neural activity for iEEG.

Our iEEG results also allowed us to ask whether temporal normalization observed in somatosensory cortex resembles the dynamics previously reported for visual cortex (Groen et al., 2022; Zhou et al., 2019). In **Supplementary Figure 1**, we plot the somatosensory iEEG time courses next to the visual iEEG time courses, enabling a direct comparison across modalities. **Supplementary Figure 2** further summarizes this comparison by visualizing the delayed normalization parameters, showing that somatosensory parameter estimates fall within the range observed across the visual hierarchy. The close correspondence in model parameters suggests that highly similar temporal dynamics operate in touch and vision. This reinforces the proposal that divisive normalization is a canonical computation shared across sensory modalities.

An alternative possibility is that the temporal nonlinearities arise from the combined outputs of a sustained and a transient channel, without the need to invoke divisive normalization. A model with a linear sustained channel and a squared transient channel has been shown to account for fMRI data in visual cortex (Horiguchi et al., 2009; Stigliani et al., 2017). Because the transient channel is squared, the two components scale differently with stimulus duration, making the model non-linear and compressive. Such a decomposition may also be physiologically motivated in somatosensation, since rapidly adapting (RA) and slowly adapting (SA) mechanoreceptor channels provide a basis for distinct transient and sustained temporal responses (Bolanowski et al., 1988; Stüttgen et al., 2006). Prior work for visual responses showed that the two-temporal-channel model fails to capture contrast-dependent changes in temporal dynamics and the temporal extent of repetition suppression (Groen et al., 2022). Nonetheless, fitting it to the present data is instructive. On the BOLD data the two-temporal-channel model performed comparably to the normalization model (cross-validated R² = 0.93 vs. 0.91) and far better than the linear model (0.67), indicating that with only the slow hemodynamic response available, fMRI data cannot distinguish the two compressive accounts (**Supplementary Figure 3**, **Figures 5A** and **6**).

The iEEG time courses resolve this ambiguity. There, the two-temporal-channel model performed no better than the linear model (cross-validated R² = 0.57 vs. 0.55) and well below delayed normalization (0.91): it did not reproduce the sustained response or the interstimulus-interval–dependent recovery from adaptation. Because its transient channel lacks history dependence, it responds identically to each pulse, and its squared transient produces spurious positive deflections at each stimulus offset that are absent from the data (**Supplementary Figure 3**). The higher temporal resolution of the iEEG measurements thus favors a history-dependent gain-control computation over a fixed two-channel decomposition, consistent with divisive normalization as a shared, canonical account across the two measurement modalities.

## Conclusion

Taken together, our results show that the temporal dynamics in human somatosensory cortex are well described by a temporal divisive normalization model, linking various nonlinear temporal signatures within a single computational framework. By combining fMRI and iEEG with matched experimental design, we demonstrated that a delayed normalization model prediction generalizes across measurement modalities. Furthermore, the similarity between somatosensory and previously reported visual temporal dynamics suggests that a common computational principle governs temporal dynamics of neural responses across modalities. Although the physical stimulus and the sensory organs differ between touch and vision, both systems encode time-varying input efficiently allowing the preservation of sensitivity to behaviorally relevant temporal structure.

## Acknowledgements

This work was supported by BRAIN Initiative Grant R01-MH111417 (to NP, JW) and NIH grant EY08266 (to MSL). We would like to thank Giovanni Piantoni at UMC Utrecht for assistance with acquiring the patient data.

## Author contributions

IMB, LL, SB, IIAG, WS, NP, MSL and JW designed research; IMB and LL collected fMRI data and analysis; IMB, LL, IIAG, NR, AF, OD, SD, WD, PD, DF were involved in the iEEG data collection and analysis; IMB, LL, MSL, and JW wrote the paper.

## 8. Supplementary Figures

**Supplementary Figure 1.**
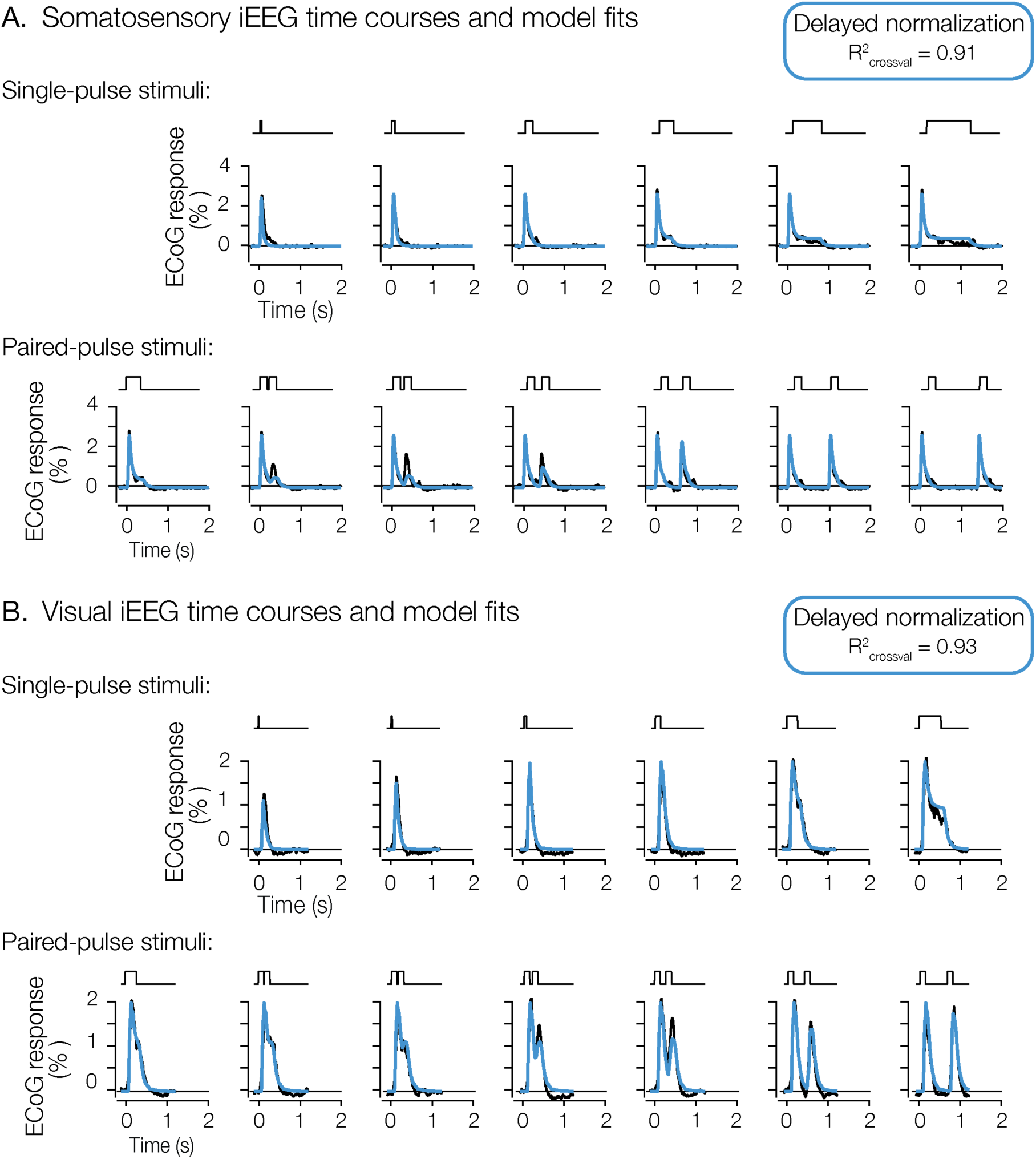
Tactile and visual iEEG time-courses. **A.** Model predictions (blue) of the iEEG responses (black) of averaged electrodes in somatosensory cortex (same as in Figure 8B). The stimulus was a single-pulse that varied in duration (first row) or a paired-pulse that varied in the inter-stimulus interval (second row). **B.** Model predictions (blue) of the iEEG responses (black) of the average of 98 electrodes in visual cortex (Groen et al., 2022). Despite different scales of stimulus durations, the tactile and visual iEEG responses are both well-captured by the same delayed normalization model. This figure is produced with show_supplementaryFigures12.m.

**Supplementary Figure 2.**
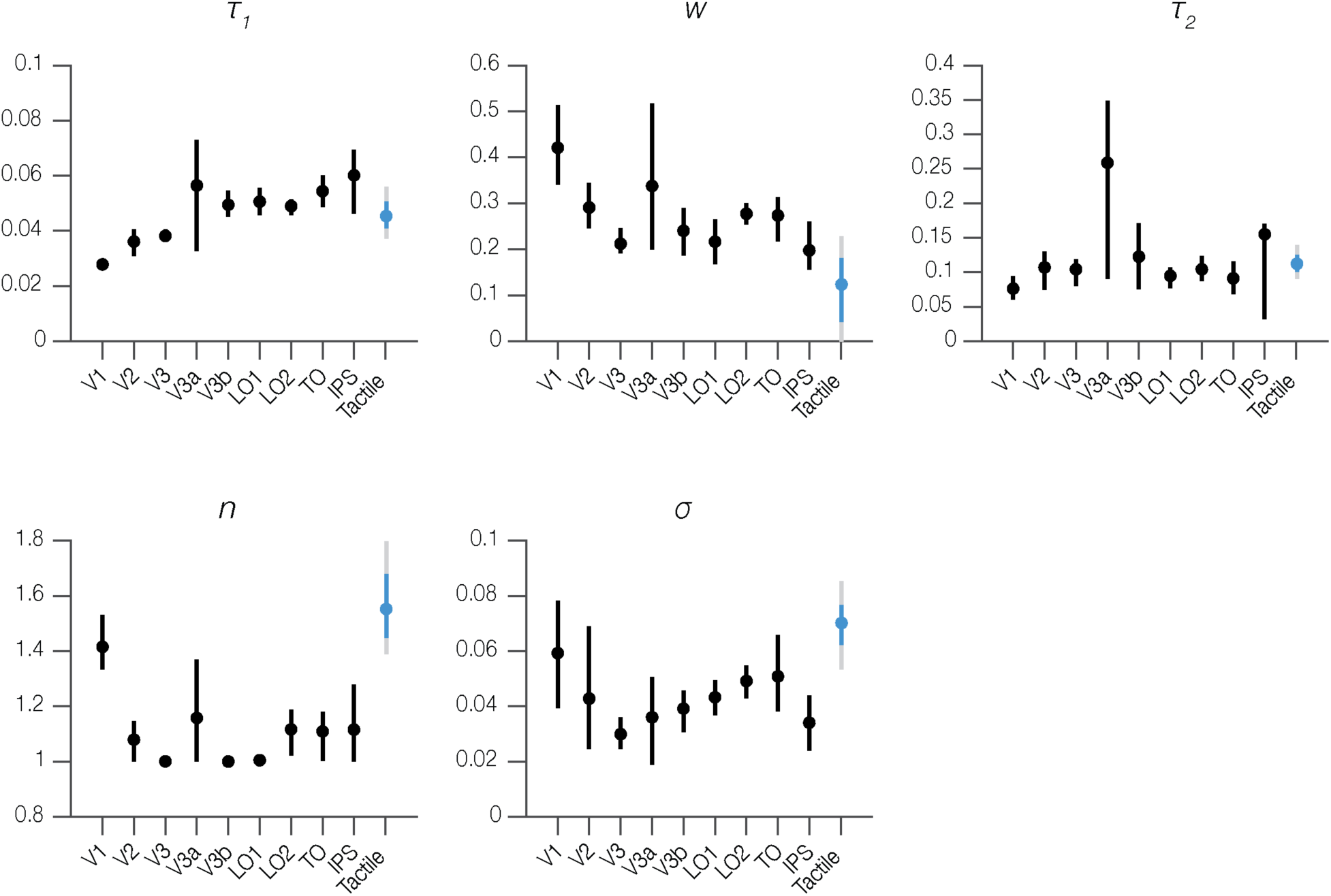
Model parameters from visual and tactile experiments. Plots show parameter estimates for nine visual areas (replotted from Groen et al., 2022) and for somatosensory cortex (replotted from Figure 8C). The tactile parameter estimates are similar to those in visual cortex and fall within the range observed across the visual hierarchy. Visual means and 68% confidence intervals are shown in black; tactile means with 68% (blue) and 95% (gray) confidence intervals. This figure is produced with show_supplementaryFigures12.m.

**Supplementary Figure 3.**
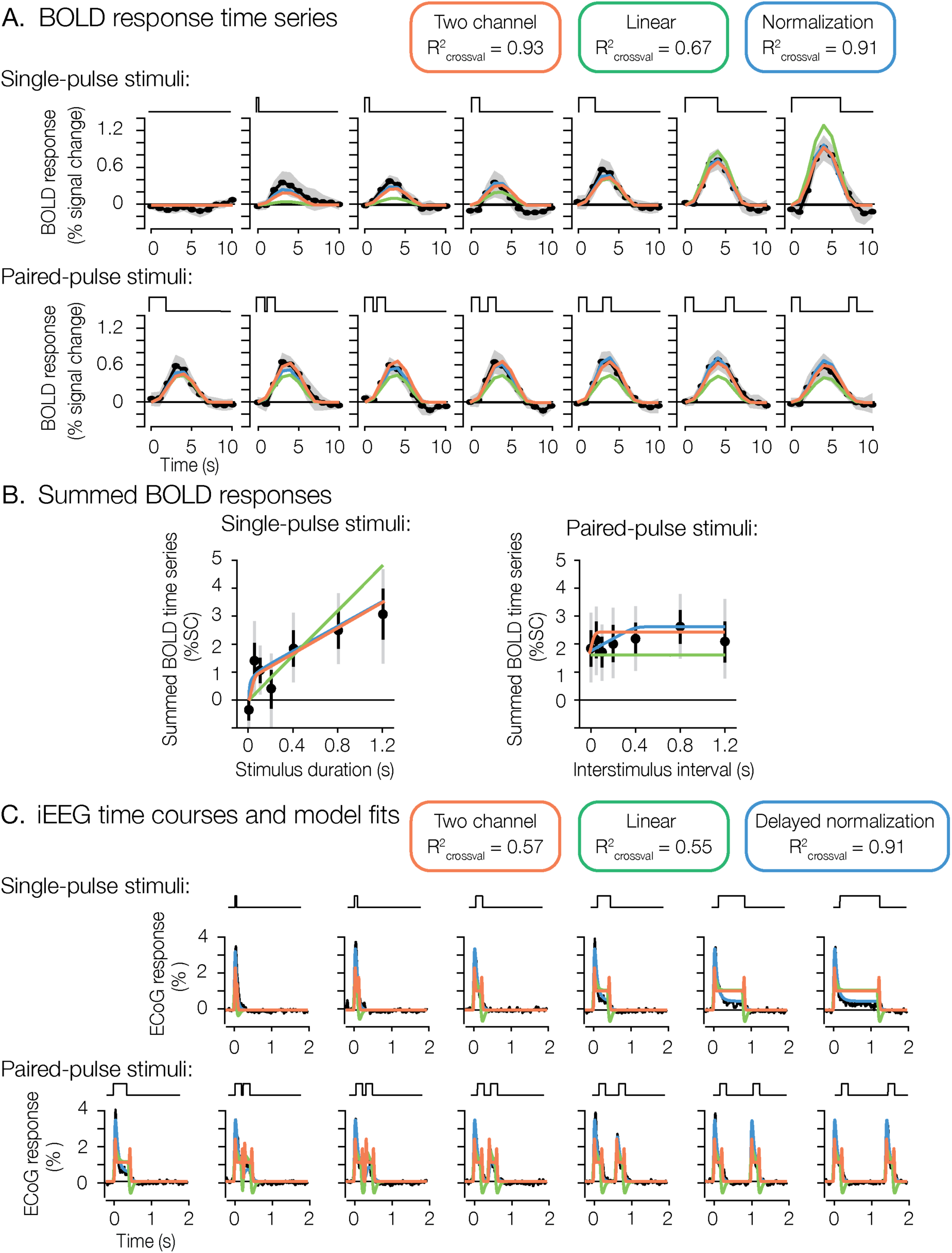
A two-temporal-channel model fit to the tactile data. The two-temporal-channel model (Stigliani et al., 2017; orange) is overlaid on the measured responses (black) alongside the linear (green) and (delayed) normalization (blue) models. **A.** fMRI single-pulse time courses (as in Figure 5A). **B.** Summed BOLD responses for single- and paired-pulse conditions (as in Figure 6). On the BOLD data the two-temporal-channel model is compressive and fits as well as the normalization model (cross-validated R² = 0.93 vs. 0.91), far exceeding the linear model (0.67); the two nonlinear models are not distinguishable at the temporal resolution of fMRI. **C.** iEEG broadband time courses (as in Figure 8B). Here, the two-temporal-channel model performs no better than the linear model (cross-validated R² = 0.57 vs. 0.55) and well below delayed normalization (0.91): it misses the sustained response and the recovery from adaptation, and its squared transient channel produces spurious positive transients at each stimulus offset (absent from the data). All cross-validated R² values are from leave-one-condition-out cross-validation. The panels were generated by show_supplementaryFigure3.m; the cross-validated R² values for all three models were recomputed in a single pipeline by recompute_allModels_CV_R2.m (leave-one-condition-out, identical held-out data).

## References

1. Acerbi, L., & Ma, W. J. (2017). Practical Bayesian Optimization for Model Fitting with Bayesian Adaptive Direct Search. Advances in Neural Information Processing Systems, 30. https://proceedings.neurips.cc/paper_files/paper/2017/hash/df0aab058ce179e4f7ab135ed4e641a9-Abstract.html

2. Andersson, J. L. R., Skare, S., & Ashburner, J. (2003). How to correct susceptibility distortions in spin-echo echo-planar images: Application to diffusion tensor imaging. NeuroImage, 20(2), 870–888. 10.1016/S1053-8119(03)00336-7

3. Arbuckle, S. A., Pruszynski, J. A., & Diedrichsen, J. (2022). Mapping the Integration of Sensory Information across Fingers in Human Sensorimotor Cortex. The Journal of Neuroscience: The Official Journal of the Society for Neuroscience, 42(26), 5173–5185. 10.1523/JNEUROSCI.2152-21.2022

4. Badde, S., Röder, B., & Heed, T. (2014). Multiple spatial representations determine touch localization on the fingers. Journal of Experimental Psychology. Human Perception and Performance, 40(2), 784–801. 10.1037/a0034690

5. Birn, R. M., & Bandettini, P. A. (2005). The effect of stimulus duty cycle and “off” duration on BOLD response linearity. NeuroImage, 27(1), 70–82. 10.1016/j.neuroimage.2005.03.040

6. Birn, R. M., Saad, Z. S., & Bandettini, P. A. (2001). Spatial Heterogeneity of the Nonlinear Dynamics in the FMRI BOLD Response. NeuroImage, 14(4), 817–826. 10.1006/nimg.2001.0873

7. Bloem, I. M., & Ling, S. (2019). Normalization governs attentional modulation within human visual cortex. Nature Communications, 10(1), 5660. 10.1038/s41467-019-13597-1

8. Bolanowski, S. J., Jr., Gescheider, G. A., Verrillo, R. T., & Checkosky, C. M. (1988). Four channels mediate the mechanical aspects of touch. The Journal of the Acoustical Society of America, 84(5), 1680–1694. 10.1121/1.397184

9. Boynton, G. M., Engel, S. A., Glover, G. H., & Heeger, D. J. (1996). Linear systems analysis of functional magnetic resonance imaging in human V1. The Journal of Neuroscience: The Official Journal of the Society for Neuroscience, 16(13), 4207–4221. 10.1523/JNEUROSCI.16-13-04207.1996

10. Boynton, G. M., Engel, S. A., & Heeger, D. J. (2012). Linear systems analysis of the fMRI signal. NeuroImage, 62(2), 975–984. 10.1016/j.neuroimage.2012.01.082

11. Brainard, D. H. (1997). The Psychophysics Toolbox. Spatial Vision, 10(4), 433–436.

12. Branco, M. P., Gaglianese, A., Glen, D. R., Hermes, D., Saad, Z. S., Petridou, N., & Ramsey, N. F. (2018). ALICE: A tool for automatic localization of intra-cranial electrodes for clinical and high-density grids. Journal of Neuroscience Methods, 301, 43–51. 10.1016/j.jneumeth.2017.10.022

13. Brands, A. M., Devore, S., Devinsky, O., Doyle, W., Flinker, A., Friedman, D., Dugan, P., Winawer, J., & Groen, I. I. A. (2024). Temporal dynamics of short-term neural adaptation across human visual cortex. PLOS Computational Biology, 20(5), e1012161. 10.1371/journal.pcbi.1012161

14. Brouwer, G. J., Arnedo, V., Offen, S., Heeger, D. J., & Grant, A. C. (2015). Normalization in human somatosensory cortex. Journal of Neurophysiology, 114(5), 2588–2599. 10.1152/jn.00939.2014

15. Carandini, M., & Heeger, D. J. (2012). Normalization as a canonical neural computation. Nature Reviews Neuroscience, 13(1), 51–62. 10.1038/nrn3136

16. Chapman, A. F., & Denison, R. N. (2025). A dynamic spatiotemporal normalization model captures perceptual and neural effects of spatial and temporal context. PLoS Biology, 23(12), e3003546. 10.1371/journal.pbio.3003546

17. Dale, A. M. (1999). Optimal experimental design for event-related fMRI. Human Brain Mapping, 8(2–3), 109–114. 10.1002/(SICI)1097-0193(1999)8:2/3<109::AID-HBM7>3.0.CO;2-W

18. Dale, A. M., Fischl, B., & Sereno, M. I. (1999). Cortical surface-based analysis. I. Segmentation and surface reconstruction. NeuroImage, 9(2), 179–194. 10.1006/nimg.1998.0395

19. de Zwart, J. A., van Gelderen, P., Jansma, J. M., Fukunaga, M., Bianciardi, M., & Duyn, J. H. (2009). Hemodynamic nonlinearities affect BOLD fMRI response timing and amplitude. NeuroImage, 47(4), 1649–1658. 10.1016/j.neuroimage.2009.06.001

20. Denison, R. N. (2024). Visual temporal attention from perception to computation. Nature Reviews Psychology, 3(4), 261–274. 10.1038/s44159-024-00294-0

21. DiCarlo, J. J., Johnson, K. O., & Hsiao, S. S. (1998). Structure of Receptive Fields in Area 3b of Primary Somatosensory Cortex in the Alert Monkey. The Journal of Neuroscience, 18(7), 2626–2645. 10.1523/JNEUROSCI.18-07-02626.1998

22. Engel, S. A., Rumelhart, D. E., Wandell, B. A., Lee, A. T., Glover, G. H., Chichilnisky, E.-J., & Shadlen, M. N. (1994). fMRI of human visual cortex. Nature, 369(6481), 525–525. 10.1038/369525a0

23. Esteban, O., Markiewicz, C. J., Blair, R. W., Moodie, C. A., Isik, A. I., Erramuzpe, A., Kent, J. D., Goncalves, M., DuPre, E., Snyder, M., Oya, H., Ghosh, S. S., Wright, J., Durnez, J., Poldrack, R. A., & Gorgolewski, K. J. (2019). fMRIPrep: A robust preprocessing pipeline for functional MRI. Nature Methods, 16(1), 111–116. 10.1038/s41592-018-0235-4

24. Fischl, B. (2004). Automatically Parcellating the Human Cerebral Cortex. Cerebral Cortex, 14(1), 11–22. 10.1093/cercor/bhg087

25. Gaglianese, A., Vansteensel, M. J., Harvey, B. M., Dumoulin, S. O., Petridou, N., & Ramsey, N. F. (2017). Correspondence between fMRI and electrophysiology during visual motion processing in human MT+. NeuroImage, 155, 480–489. 10.1016/j.neuroimage.2017.04.007

26. Glasser, M. F., Coalson, T. S., Robinson, E. C., Hacker, C. D., Harwell, J., Yacoub, E., Ugurbil, K., Andersson, J., Beckmann, C. F., Jenkinson, M., Smith, S. M., & Van Essen, D. C. (2016). A multi-modal parcellation of human cerebral cortex. Nature, 536(7615), 171–178. 10.1038/nature18933

27. Glover, G. H. (1999). Deconvolution of Impulse Response in Event-Related BOLD fMRI1. NeuroImage, 9(4), 416–429. 10.1006/nimg.1998.0419

28. Gorgolewski, K., Burns, C. D., Madison, C., Clark, D., Halchenko, Y. O., Waskom, M. L., & Ghosh, S. S. (2011). Nipype: A flexible, lightweight and extensible neuroimaging data processing framework in python. Frontiers in Neuroinformatics, 5, 13. 10.3389/fninf.2011.00013

29. Greve, D. N., & Fischl, B. (2009). Accurate and robust brain image alignment using boundary-based registration. NeuroImage, 48(1), 63–72. 10.1016/j.neuroimage.2009.06.060

30. Groen, I. I. A., Piantoni, G., Montenegro, S., Flinker, A., Devore, S., Devinsky, O., Doyle, W., Dugan, P., Friedman, D., Ramsey, N. F., Petridou, N., & Winawer, J. (2022). Temporal Dynamics of Neural Responses in Human Visual Cortex. Journal of Neuroscience, 42(40), 7562–7580. 10.1523/JNEUROSCI.1812-21.2022

31. Heeger, D. J. (1992). Normalization of cell responses in cat striate cortex. Visual Neuroscience, 9(2), 181–197. 10.1017/S0952523800009640

32. Holdgraf, C., Appelhoff, S., Bickel, S., Bouchard, K., D’Ambrosio, S., David, O., Devinsky, O., Dichter, B., Flinker, A., Foster, B. L., Gorgolewski, K. J., Groen, I., Groppe, D., Gunduz, A., Hamilton, L., Honey, C. J., Jas, M., Knight, R., Lachaux, J.-P., … Hermes, D. (2019). iEEG-BIDS, extending the Brain Imaging Data Structure specification to human intracranial electrophysiology. Scientific Data, 6(1), 102. 10.1038/s41597-019-0105-7

33. Horiguchi, H., Nakadomari, S., Misaki, M., & Wandell, B. A. (2009). Two temporal channels in human V1 identified using fMRI. NeuroImage, 47(1), 273–280. 10.1016/j.neuroimage.2009.03.078

34. Hyvärinen, J., & Poranen, A. (1978). Receptive field integration and submodality convergence in the hand area of the post-central gyrus of the alert monkey. The Journal of Physiology, 283, 539–556. 10.1113/jphysiol.1978.sp012518

35. Jenkinson, M., Bannister, P., Brady, M., & Smith, S. (2002). Improved Optimization for the Robust and Accurate Linear Registration and Motion Correction of Brain Images. NeuroImage, 17(2), 825–841. 10.1006/nimg.2002.1132

36. Kaas, J. H. (1993). The functional organization of somatosensory cortex in primates. Annals of Anatomy - Anatomischer Anzeiger, 175(6), 509–518. 10.1016/S0940-9602(11)80212-8

37. Kay, K. N., Rokem, A., Winawer, J., Dougherty, R. F., & Wandell, B. A. (2013). GLMdenoise: A fast, automated technique for denoising task-based fMRI data. Frontiers in Neuroscience, 7, 247. 10.3389/fnins.2013.00247

38. Keung, W., Hagen, T. A., & Wilson, R. C. (2020). A divisive model of evidence accumulation explains uneven weighting of evidence over time. Nature Communications, 11(1), 2160. 10.1038/s41467-020-15630-0

39. Kupers, E. R., Kim, I., & Grill-Spector, K. (2024). Rethinking simultaneous suppression in visual cortex via compressive spatiotemporal population receptive fields. Nature Communications, 15(1), 6885. 10.1038/s41467-024-51243-7

40. Lee, J., & Maunsell, J. H. R. (2009). A Normalization Model of Attentional Modulation of Single Unit Responses. PLOS ONE, 4(2), e4651. 10.1371/journal.pone.0004651

41. Lewis, L. D., Setsompop, K., Rosen, B. R., & Polimeni, J. R. (2018). Stimulus-dependent hemodynamic response timing across the human subcortical-cortical visual pathway identified through high spatiotemporal resolution 7T fMRI. NeuroImage, 181, 279–291. 10.1016/j.neuroimage.2018.06.056

42. Lisberger, S. G., & Movshon, J. A. (1999). Visual Motion Analysis for Pursuit Eye Movements in Area MT of Macaque Monkeys. Journal of Neuroscience, 19(6), 2224–2246. 10.1523/JNEUROSCI.19-06-02224.1999

43. Louie, K., Khaw, M. W., & Glimcher, P. W. (2013). Normalization is a general neural mechanism for context-dependent decision making. Proceedings of the National Academy of Sciences of the United States of America, 110(15), 6139–6144. 10.1073/pnas.1217854110

44. Mikaelian, S., & Simoncelli, E. P. (2001). Modeling temporal response characteristics of V1 neurons with a dynamic normalization model. Neurocomputing, Computational Neuroscience: Trends in Research 2001, 38-40, 1461–1467. 10.1016/S0925-2312(01)00529-X

45. Miller, K. J., Honey, C. J., Hermes, D., Rao, R. P., denNijs, M., & Ojemann, J. G. (2014). Broadband changes in the cortical surface potential track activation of functionally diverse neuronal populations. NeuroImage, New Horizons for Neural Oscillations, 85, 711–720. 10.1016/j.neuroimage.2013.08.070

46. Olsen, S. R., Bhandawat, V., & Wilson, R. I. (2010). Divisive Normalization in Olfactory Population Codes. Neuron, 66(2), 287–299. 10.1016/j.neuron.2010.04.009

47. Oostenveld, R., Fries, P., Maris, E., & Schoffelen, J.-M. (2011). FieldTrip: Open source software for advanced analysis of MEG, EEG, and invasive electrophysiological data. Computational Intelligence and Neuroscience, 2011, 156869. 10.1155/2011/156869

48. Pelli, D. G. (1997). The VideoToolbox software for visual psychophysics: Transforming numbers into movies. Spatial Vision, 10(4), 437–442.

49. Penfield, W., & Rasmussen, T. (1950). The cerebral cortex of man; a clinical study of localization of function (pp. xv, 248). Macmillan.

50. Pfeuffer, J., McCullough, J. C., Van de Moortele, P.-F., Ugurbil, K., & Hu, X. (2003). Spatial dependence of the nonlinear BOLD response at short stimulus duration. NeuroImage, 18(4), 990–1000. 10.1016/S1053-8119(03)00035-1

51. Polimeni, J. R., & Lewis, L. D. (2021). Imaging faster neural dynamics with fast fMRI: A need for updated models of the hemodynamic response. Progress in Neurobiology, How High Spatiotemporal Resolution fMRI Can Advance Neuroscience, 207, 102174. 10.1016/j.pneurobio.2021.102174

52. Reed, J. L., Pouget, P., Qi, H.-X., Zhou, Z., Bernard, M. R., Burish, M. J., Haitas, J., Bonds, A. B., & Kaas, J. H. (2008). Widespread spatial integration in primary somatosensory cortex. Proceedings of the National Academy of Sciences of the United States of America, 105(29), 10233–10237. 10.1073/pnas.0803800105

53. Reynolds, J. H., & Heeger, D. J. (2009). The Normalization Model of Attention. Neuron, 61(2), 168–185. 10.1016/j.neuron.2009.01.002

54. Sanchez-Panchuelo, R. M., Francis, S., Bowtell, R., & Schluppeck, D. (2010). Mapping human somatosensory cortex in individual subjects with 7T functional MRI. Journal of Neurophysiology, 103(5), 2544–2556. 10.1152/jn.01017.2009

55. Schellekens, W., Thio, M., Badde, S., Winawer, J., Ramsey, N., & Petridou, N. (2021). A touch of hierarchy: Population receptive fields reveal fingertip integration in Brodmann areas in human primary somatosensory cortex. Brain Structure & Function, 226(7), 2099–2112. 10.1007/s00429-021-02309-5

56. Sereno, M. I., Dale, A. M., Reppas, J. B., Kwong, K. K., Belliveau, J. W., Brady, T. J., Rosen, B. R., & Tootell, R. B. (1995). Borders of multiple visual areas in humans revealed by functional magnetic resonance imaging. Science, 268(5212), 889–893. 10.1126/science.7754376

57. Siero, J. C., Hermes, D., Hoogduin, H., Luijten, P. R., Petridou, N., & Ramsey, N. F. (2013). BOLD consistently matches electrophysiology in human sensorimotor cortex at increasing movement rates: A combined 7T fMRI and ECoG study on neurovascular coupling. Journal of Cerebral Blood Flow & Metabolism, 33(9), 1448–1456. 10.1038/jcbfm.2013.97

58. Stigliani, A., Jeska, B., & Grill-Spector, K. (2017). Encoding model of temporal processing in human visual cortex. Proceedings of the National Academy of Sciences, 114(51), E11047–E11056. 10.1073/pnas.1704877114

59. Stigliani, A., Jeska, B., & Grill-Spector, K. (2019). Differential sustained and transient temporal processing across visual streams. PLoS Computational Biology, 15(5), e1007011. 10.1371/journal.pcbi.1007011

60. Stüttgen, M. C., Rüter, J., & Schwarz, C. (2006). Two Psychophysical Channels of Whisker Deflection in Rats Align with Two Neuronal Classes of Primary Afferents. Journal of Neuroscience, 26(30), 7933–7941. 10.1523/JNEUROSCI.1864-06.2006

61. Tolhurst, D. J., Walker, N. S., Thompson, I. D., & Dean, A. F. (1980). Non-linearities of temporal summation in neurones in area 17 of the cat. Experimental Brain Research, 38(4), 431–435. 10.1007/BF00237523

62. Tommerdahl, M., Favorov, O. V., & Whitsel, B. L. (2010). Dynamic representations of the somatosensory cortex. Neuroscience & Biobehavioral Reviews, Touch, Temperature, Pain/Itch and Pleasure, 34(2), 160–170. 10.1016/j.neubiorev.2009.08.009

63. Tustison, N. J., Avants, B. B., Cook, P. A., Zheng, Y., Egan, A., Yushkevich, P. A., & Gee, J. C. (2010). N4ITK: Improved N3 bias correction. IEEE Transactions on Medical Imaging, 29(6), 1310–1320. 10.1109/TMI.2010.2046908

64. Willmore, B. D. B., & King, A. J. (2023). Adaptation in auditory processing. Physiological Reviews, 103(2), 1025–1058. 10.1152/physrev.00011.2022

65. Yeşilyurt, B., Uğurbil, K., & Uludağ, K. (2008). Dynamics and nonlinearities of the BOLD response at very short stimulus durations. Magnetic Resonance Imaging, 26(7), 853–862. 10.1016/j.mri.2008.01.008

66. Zhang, N., Zhu, X.-H., & Chen, W. (2008). Investigating the source of BOLD nonlinearity in human visual cortex in response to paired visual stimuli. NeuroImage, 43(2), 204–212. 10.1016/j.neuroimage.2008.06.033

67. Zhang, Y., Brady, M., & Smith, S. (2001). Segmentation of brain MR images through a hidden Markov random field model and the expectation-maximization algorithm. IEEE Transactions on Medical Imaging, 20(1), 45–57. 10.1109/42.906424

68. Zhou, J., Benson, N. C., Kay, K. N., & Winawer, J. (2018). Compressive Temporal Summation in Human Visual Cortex. Journal of Neuroscience, 38(3), 691–709. 10.1523/JNEUROSCI.1724-17.2017

69. Zhou, J., Benson, N. C., Kay, K., & Winawer, J. (2019). Predicting neuronal dynamics with a delayed gain control model. PLOS Computational Biology, 15(11), e1007484. 10.1371/journal.pcbi.1007484

