## Supplementary material for "Normalization accounts for temporal dynamics in human somatosensory cortex": Figure S1, Figure S2 and Figure S3

### 8. Supplementary Figures

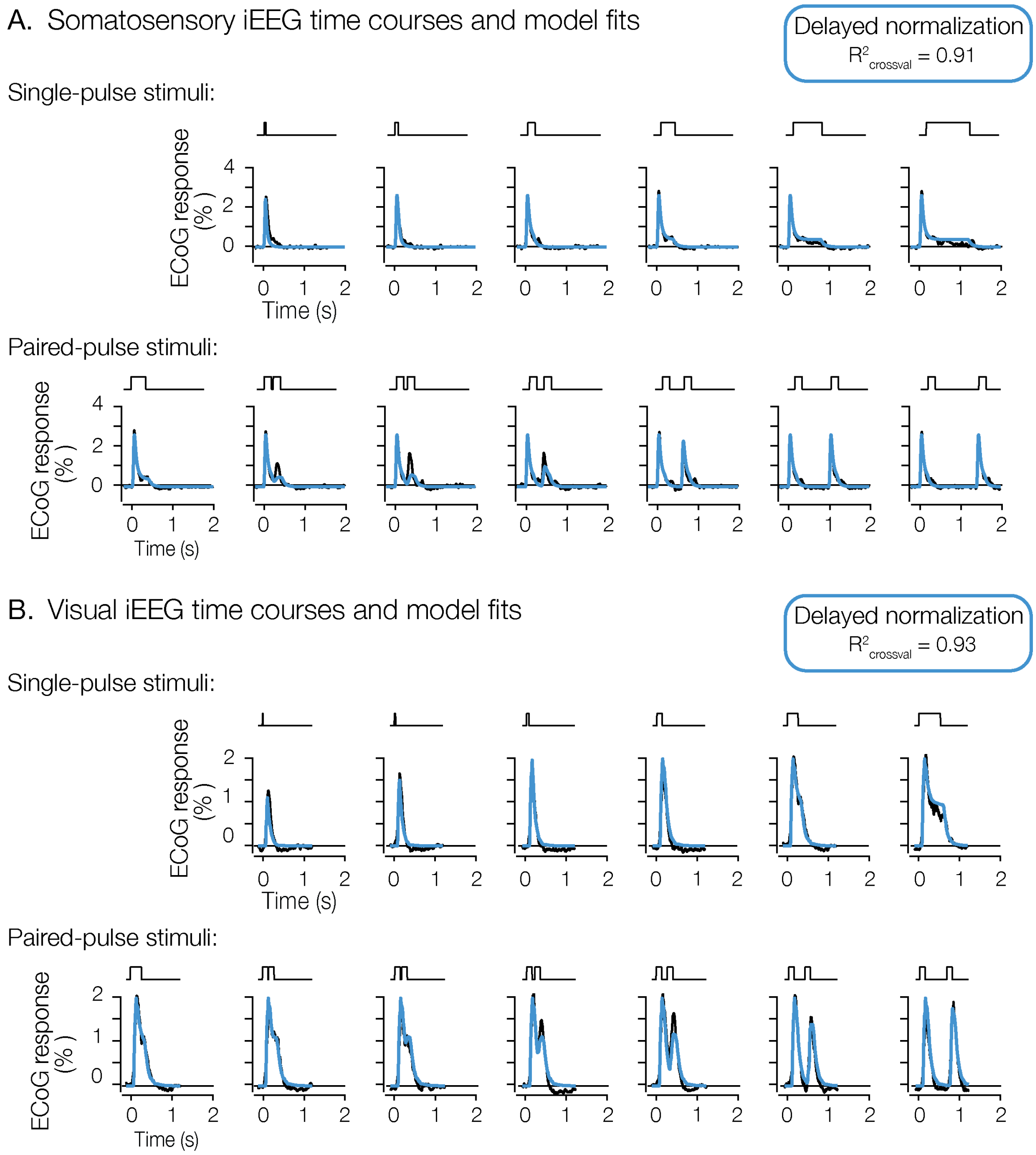


**Supplementary Figure 1. Tactile and visual iEEG time-courses. A.** Model predictions (blue) of the iEEG responses (black) of averaged electrodes in somatosensory cortex (same as in Figure 8B). The stimulus was a single-pulse that varied in duration (first row) or a paired-pulse that varied in the inter-stimulus interval (second row). **B.** Model predictions (blue) of the iEEG responses (black) of the average of 98 electrodes in visual cortex (Groen et al., 2022). Despite different scales of stimulus durations, the tactile and visual iEEG responses are both well-captured by the same delayed normalization model. This figure is produced with show_supplementaryFigures12.m.

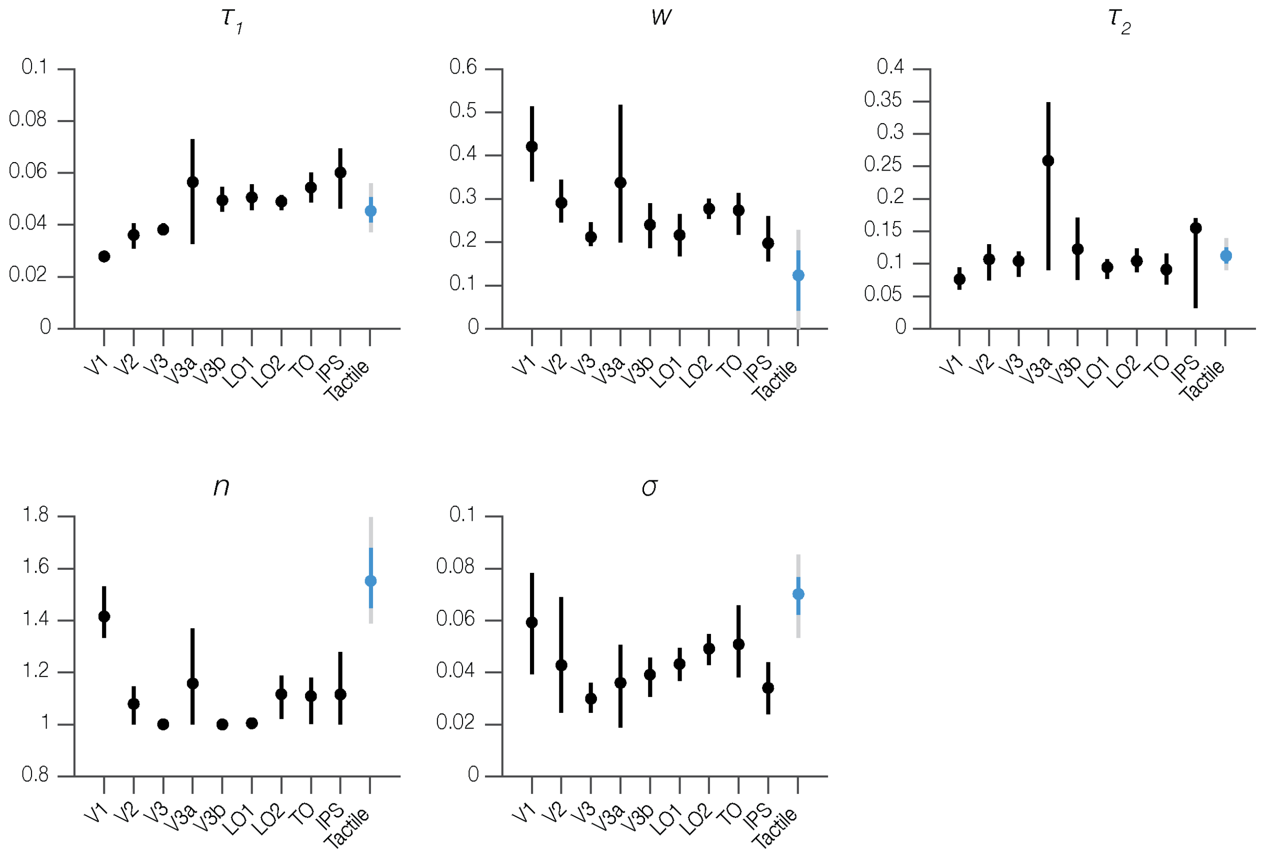


**Supplementary Figure 2. Model parameters from visual and tactile experiments.** Plots show parameter estimates for nine visual areas (replotted from Groen et al., 2022) and for somatosensory cortex (replotted from Figure 8C). The tactile parameter estimates are similar to those in visual cortex and fall within the range observed across the visual hierarchy. Visual means and 68% confidence intervals are shown in black; tactile means with 68% (blue) and 95% (gray) confidence intervals. This figure is produced with show_supplementaryFigures12.m.

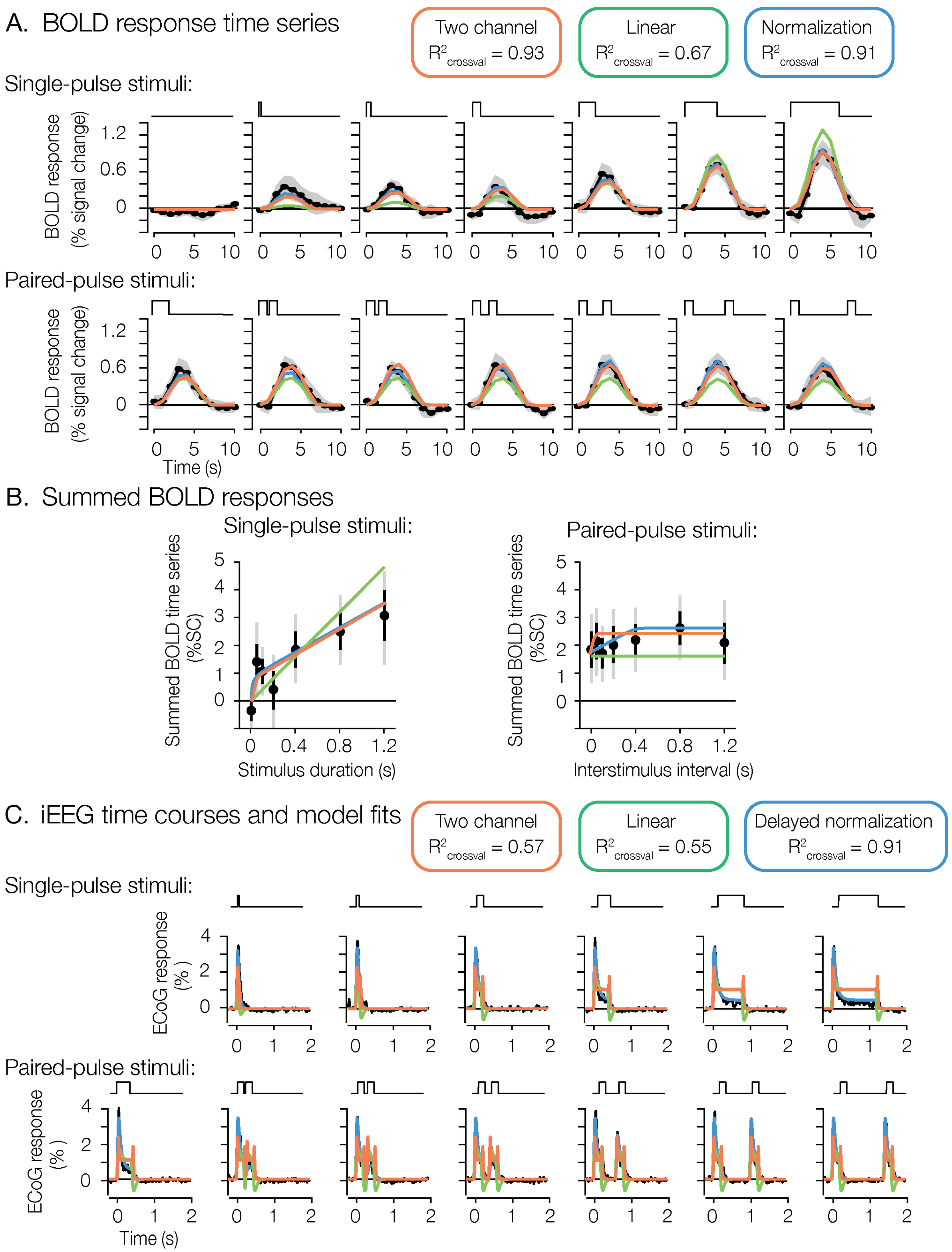


**Supplementary Figure 3. A two-temporal-channel model fit to the tactile data.** The two-temporal-channel model (Stigliani et al., 2017; orange) is overlaid on the measured responses (black) alongside the linear (green) and (delayed) normalization (blue) models. **A.** fMRI single-pulse time courses (as in Figure 5A). **B.** Summed BOLD responses for single- and paired-pulse conditions (as in Figure 6). On the BOLD data the two-temporal-channel model is compressive and fits as well as the normalization model (cross-validated $R^2$ = 0.93 vs. 0.91), far exceeding the linear model (0.67); the two nonlinear models are not distinguishable at the temporal resolution of fMRI. **C.** iEEG broadband time courses (as in Figure 8B). Here, the two-temporal-channel model performs no better than the linear model (cross-validated $R^2$ = 0.57 vs. 0.55) and well below delayed normalization (0.91): it misses the sustained response and the recovery from adaptation, and its squared transient channel produces spurious positive transients at each stimulus offset (absent from the data). All cross-validated $R^2$ values are from leave-one-condition-out cross-validation. The panels were generated by show_supplementaryFigure3.m; the cross-validated $R^2$ values for all three models were recomputed in a single pipeline by recompute_allModels_CV_R2.m (leave-one-condition-out, identical held-out data).
